# Combinatorial phenotypic landscape enables bacterial resistance to phage infection

**DOI:** 10.1101/2025.01.13.632860

**Authors:** Anika Gupta, Norma Morella, Dmitry Sutormin, Naisi Li, Karl Gaisser, Alexander Robertson, Yaroslav Ispolatov, Georg Seelig, Neelendu Dey, Anna Kuchina

## Abstract

Success of phage therapies is limited by bacterial defenses against phages. While a large variety of anti-phage defense mechanisms has been characterized, how expression of these systems is distributed across individual cells and how their combined activities translate into protection from phages has not been studied. Using bacterial single-cell RNA sequencing, we profiled the transcriptomes of ∼50,000 cells from cultures of a human pathobiont, *Bacteroides fragilis,* infected with a lytic bacteriophage. We quantified the asynchronous progression of phage infection in single bacterial cells and reconstructed the infection timeline, characterizing both host and phage transcriptomic changes as infection unfolded. We discovered a subpopulation of bacteria that remained uninfected and determined the heterogeneously expressed host factors associated with protection. Each cell’s vulnerability to phage infection was defined by combinatorial phase-variable expression of multiple genetic loci, including capsular polysaccharide (CPS) biosynthesis pathways, restriction-modification systems (RM), and a previously uncharacterized operon likely encoding fimbrial genes. By acting together, these heterogeneously expressed phase-variable systems and anti-phage defense mechanisms create a phenotypic landscape where distinct protective combinations enable the survival and re-growth of bacteria expressing these phenotypes without acquiring additional mutations. The emerging model of complementary action of multiple protective mechanisms heterogeneously expressed across an isogenic bacterial population showcases the potent role of phase variation and stochasticity in bacterial anti-phage defenses.

**One Sentence Summary:** Combinatorial phenotypic states with differential vulnerability to phage infection across a *Bacteroides fragilis* population enable a small number of super-resistant bacterial cells to evade the phage without the need for acquiring mutations.

## Main Text

The human gut microbiome contains multiple species of bacteriophages (hereafter “phages”) which interact with specific bacterial hosts(*1*). Since lytic phage infections result in rapid host death, phages act as potent

modulators of microbiome composition and potential tools for combating drug-resistant infections. Phage therapy, in the form of application of a single bacteriophage or a ‘phage cocktail’ to treat an infection, has been used in clinical trials against enteric pathogens(*2–4*). However, the emergence of resistance limits current therapeutic development and application of phages.

Bacteria employ multiple resistance mechanisms to defend themselves, including barriers to phage entry, such as polysaccharide capsules, and an expanding arsenal of internal systems limiting phage infection. Beside well-known nucleic-acid degrading antiphage defense mechanisms such as restriction-modification (RM) systems and CRISPR-Cas, over a hundred bacterial immunity systems are now known in a range of diverse species(*5*). Despite these advances, relatively few of these systems have been investigated in their natural host, with the majority of them studied via heterologous expression in a model *E. coli* strain(*6*). In this model, high constitutive induction of these systems does not reflect their natural expression patterns, which frequently vary across individual bacterial cells even in pure cultures under identical conditions(*7*, *8*). This behavior, termed phenotypic heterogeneity, is assumed to enhance the population-level fitness of the bacterium, for example, when changes in the environment confer a strong benefit upon specific phenotypes. The bacterial species inhabiting the human gut are known to exhibit extensive phenotypic heterogeneity contributing to defenses against phages. One example is the phase-variable expression of capsular polysaccharide (CPS) genes in *Bacteroides* spp., mediated by the stochastic inversion of promoters controlling the expression of genes to produce each of the multiple CPS types(*9*, *10*), some of which provide protection against phages(*11*). The heterogeneity of expression of other types of anti-phage defense systems such as CRISPR-Cas is also becoming evident(*12*).

Many bacterial species harbor multiple anti-phage defense systems in their genome with potentially synergistic contributions to protection from phages(*6*, *13*). However, interactions between most anti-phage protective mechanisms are poorly understood. Additionally, the heterogeneity of these systems’ expression in individual cells creates combinatorial phenotypic states across the population with distinct vulnerabilities to attacks by different phages. Survival of rare phenotypically protected cells may then provide an opportunity for the development of genetic resistance. Due to the challenges of single-cell gene expression profiling in human gut microbiota members, the combinatorial contributions of heterogeneously expressed defense systems on phage infection outcomes in these hosts have not been explored.

Here, we use microbial split-pool ligation transcriptomics (microSPLiT), a bacterial single-cell RNA sequencing technology, to explore the progression of phage infection across individual host cells and identify phase-variable regions combinatorially expressed in a subpopulation of cells that is phenotypically protected from infection, giving rise to phage resistance. Specifically, we profile gene expression in individual cells of *Bacteroides fragilis,* a commensal gut bacterium and pathobiont implicated in multiple disease states such as colorectal cancer(*14*, *15*), abscesses(*16*, *17*), undernutrition(*18*), and inflammatory bowel disease(*19*), infected with a lytic bacteriophage isolated from human waste.

## Single-cell RNA sequencing identifies phage-infected and -uninfected bacterial subpopulations

We isolated a lytic bacteriophage from King County wastewater that we named *Bacteroides* phage Bf12P1, with species- and strain-level host specificity to *Bacteroides fragilis* NCTC 9343. Whole genome sequencing (WGS) and morphological characterization (**Figure 1A**, **Supplementary** Figure 1A) were used to identify Bf12P1 as a novel siphovirus. Bf12P1 has a 47,370 bp-genome with 72 coding sequences and is most closely related to *Bacteroides* phage Barc2635 (GenBank: MN078104.1) with a nucleotide similarity of 89.0% (**Figure 1B**, **Supplementary** Figure 1B) and viral sequences obtained from human gut metagenomes. Bf12P1 readily infects and lyses its host within 2 hours, as determined through liquid growth curves and visible lysis on plates (**Supplementary** Figure 1C).

**Fig. 1:**
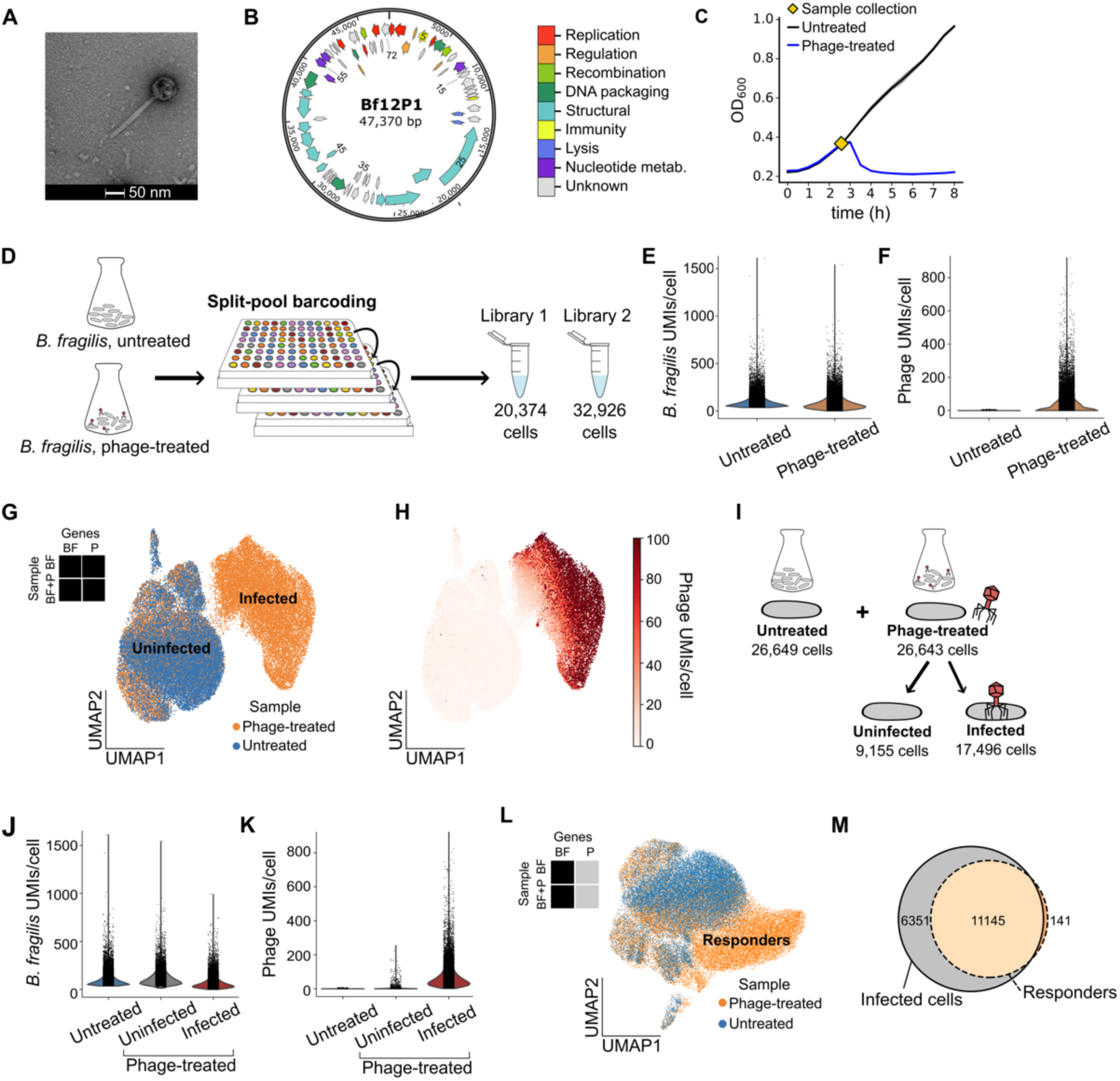
Single-cell transcriptomics of *B. fragilis* treated with Bf12P1 phage detects infected and uninfected cells within a single sample. **(A)** Electron microscopy image of siphovirus Bf12P1. **(B)** Bf12P1 genome with feature colors corresponding to putative gene functions and numerical labels indicating locations of each gene. **(C)** Growth curves of untreated and phage-treated *B. fragilis* from an experiment prior to microSPLiT sampling (MOI of 0.00004). Symbol marks the time of sample collection chosen for the future M30 and F3 experiments. **(D)** Summary of F3 microSPLiT experiment on both untreated and phage-treated *B. fragilis* samples producing two replicate sequencing libraries. **(E)** *B. fragilis,* and **(F)** Bf12P1 UMIs/cell for untreated and phage-treated samples (F3 experiment, combined replicates). **(G)** UMAP embedding of both treated and untreated samples, using both *B. fragilis* and Bf12P1 genes, colored by sample. The two distinct parts of the UMAP plot representing the infected or uninfected cells are labeled accordingly. **(H)** Phage UMIs/cell overlaid on the UMAP from panel G. **(I)** Sample definitions. The phage-treated sample after sequencing is divided into uninfected and infected subpopulations based on the location on the UMAP plot with indicated cell numbers. **(J and K)** *B. fragilis* and Bf12P1 UMIs/cell, respectively, for all subpopulations of cells including uninfected and infected cells. **(L)** UMAP embedding of both phage- treated and untreated samples constructed using only *B. fragilis* transcripts, colored by sample. A distinct cluster of cells from the phage-treated sample is labeled “responders.” **(M)** Venn diagram showing the overlap between cells in the “responders” cluster with the cells in the infected subpopulation of cells defined in panel G.

To investigate the differences in infection susceptibility across individual cells in a population of *B. fragilis*, we infected the sample at multiplicity of infection (MOI) 0.00004 and collected phage-treated cells after 2.5h of infection (“Phage-treated”), just before the projected onset of phage-induced lysis. In parallel, a set of *B. fragilis* cultures was prepared from the same stock to which phage had not been added (“Untreated”), and samples were collected at the same time point and optical density (**Figure 1C**). Finally, we collected a similar set of samples using media with added deoxycholic acid (DCA), a secondary bile acid common in the gut environment(*20*). We immediately fixed the samples with formaldehyde and proceeded with microSPLiT, a method we previously developed for single-cell RNA sequencing of bacteria(*8*).

We performed two independent microSPLiT experiments, named “M30” and “F3”. While yielding similar results, F3 retrieved substantially more cells with high transcript content due to workflow optimizations (**Supplementary Note 1, Supplementary** Figures 2-3, Materials and Methods). After filtering out low quality cells, the no-DCA dataset contained 53,300 cells from two replicate F3 libraries (**Figure 1D**), with medians of 79 and 66 *B. fragilis* mRNA transcripts per cell for the untreated and phage-treated samples, respectively (**Figure 1E**). We also recovered 27 median phage transcripts per cell for the phage-treated sample (**Figure 1F**). The summed gene expression profile correlated with the bulk RNA sequencing performed earlier (**Supplementary** Figure 4). Unsupervised clustering of both samples revealed two main subpopulations of cells, largely discriminated by the presence of the phage transcripts (**Figure 1G-H and Supplementary** Figure 3A**, C**). Approximately two-thirds of the cells from the phage-treated sample contained phage transcripts and clustered separately from the untreated sample. The remaining phage-treated cells clustered together with untreated cells and lacked phage transcripts. We named this subpopulation of cells from the phage-treated sample “uninfected,” as opposed to the “infected” subpopulation of the same sample (**Figure 1I**). The infected subpopulation displayed a reduced median of 50 *B. fragilis* mRNA UMIs/cell, whereas we retrieved 100 median mRNA UMIs/cell for the uninfected subpopulation (**Figure 1J**). The Bf12P1 genes were detected at a median of 51 UMIs/cell for the infected subpopulation (**Figure 1K**). A similar distinction was observed in samples exposed to DCA (**Supplementary** Figure 5A).

Clustering on the basis of both host and phage gene expression could be primarily driven by the presence of phage reads. To reveal host-specific transcriptional changes in response to bacteriophage infection, we removed phage transcripts from our dataset and re-clustered the data (**Figure 1L**). We observed a new cluster, populated only by cells from the phage-treated sample, that we assumed was exhibiting the transcriptional response to phage infection; thus, we named it “responders” (**Figure 1L**). Indeed, when we extracted the barcodes of cells in the “responders” cluster and retrieved phage transcripts associated with these cellular identifiers, the vast majority of cells (98.7%) had an associated phage load and were previously assigned to the infected subpopulation (**Figure 1M**). This analysis confirmed that the majority of infected *B. fragilis* cells displayed a distinct host gene expression profile associated with phage infection.

Examining the transcriptomic profile in the “responders” cluster, we found that it was predominantly defined by the expression of RNA polymerase subunits, replication protein genes, and multiple ribonucleoside reductases (**Supplementary Note 2**). While overall this gene expression pattern reflected the expected host response to phage infection, it was an average representation of cells at different points in the infectious cycle. Prior to sampling, the phage-treated cell population had likely undergone multiple infectious cycles, the timing of which presumably varied between individual cells due to the intrinsic heterogeneity of host-phage interactions on the single cell level. Thus, we next turned to resolving cell-to-cell variability in the progression of phage infection within the infected cells.

## Single-cell host and phage gene expression reveals infection timeline and uncovers early and late phage gene modules

Subclustering the phage-treated sample by itself unveiled a gradient of phage transcript density across cells with a clear separation between the infected and uninfected subpopulations **(Figure 2A**). To learn whether this pattern reflected differences in infection development between cells, we computed a pseudotime trajectory of the phage-treated sample, reconstructing the ordering of cells along a biological process, which aligned well with the distribution of phage transcripts across cells (**Figure 2B**). The relative position of a cell within the pseudotime trajectory is indicated by a numerical value between 0 and 1. While *B. fragilis* transcript content declined with pseudotime, reaching almost negligible values above a pseudotime value of 0.7, phage transcripts increased concurrently (**Figure 2C**). We detected minimal to no phage transcripts between values 0.0 to 0.2, as cells within this range were primarily uninfected. These transcriptomic patterns show that the calculated pseudotime trajectory reflects the progression of phage infection within *B. fragilis* cells, with an initial rapid shutdown of most host gene expression followed by induction of cellular machinery for phage replication, and degradation of the host, accompanied by an increase in phage abundance.

**Fig. 2:**
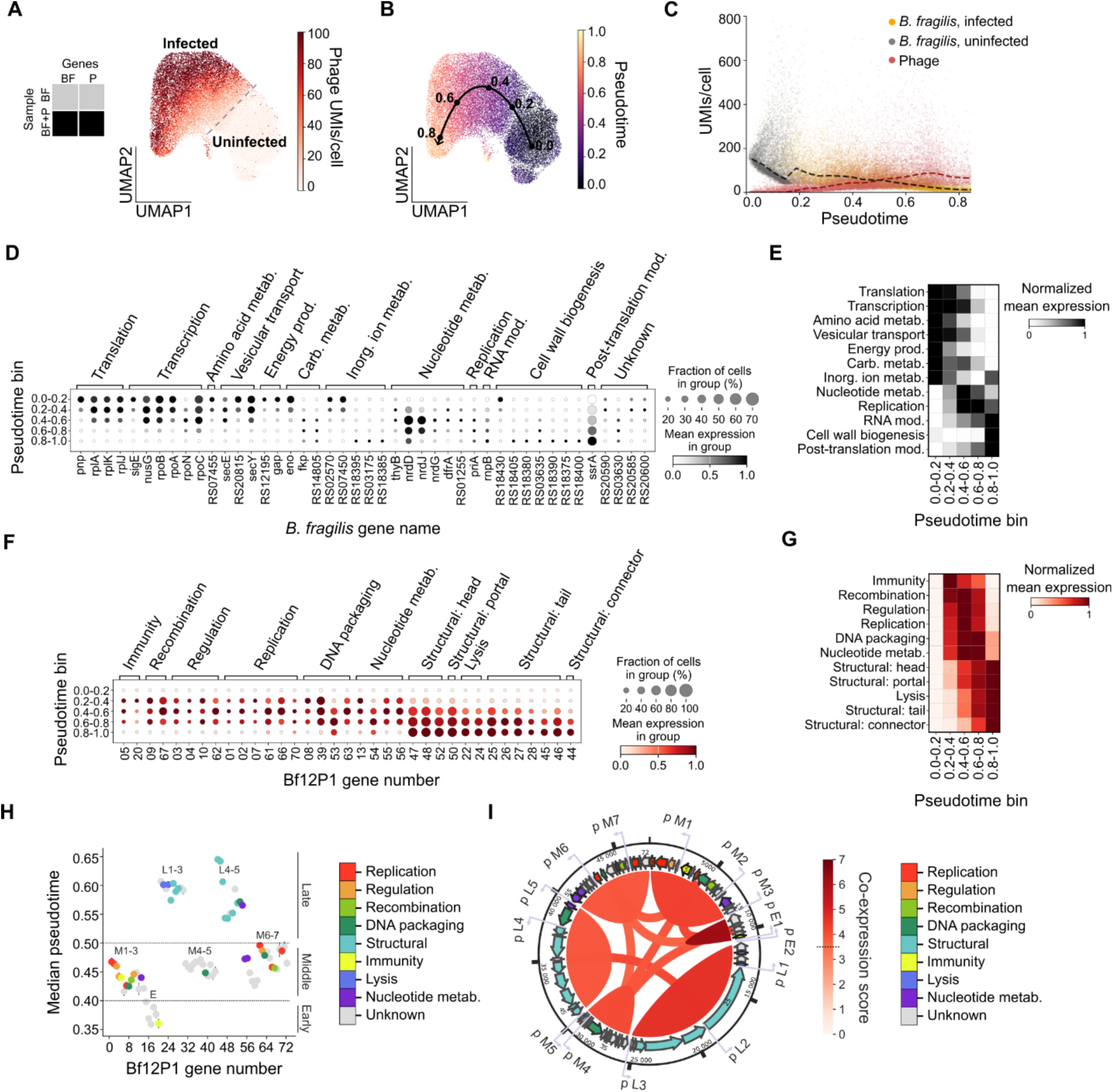
Host and phage gene expression profiles along the calculated infection trajectory. **(A)** UMAP embedding of the phage-treated sample using both *B. fragilis* and Bf12P1 genes, divided between uninfected and infected cell subpopulations. Phage UMIs/cell are shown in red. **(B)** UMAP plot from panel A colored by calculated diffusion pseudotime values, overlaid with a trajectory connecting cell regions of approximately 0.0, 0.2, 0.4, 0.6, and 0.8 pseudotime values. **(C)** *B. fragilis* UMIs/cell (gray and yellow for uninfected and infected cells, respectively) and phage UMIs/cell (red) in the phage-treated sample plotted against the pseudotime values with each dot representing a single cell. Median UMIs/cell across pseudotime are shown as a black line for *B. fragilis* and a red line for phage. **(D)** Expression of top 10 differentially expressed genes in the phage-treated sample across 5 equally sized pseudotime bins, categorized by function. **(E)** Summed raw *B. fragilis* gene expression for each functional category in each pseudotime bin, normalized by bin. **(F)** Expression of all phage genes in the phage-treated sample across 5 pseudotime bins, grouped according to function. **(G)** Summed raw Bf12P1 gene expression for each functional category in each pseudotime bin, normalized by bin. **(H)** Median pseudotime of expression for each phage gene shown as a dot colored according to functional category and arranged by genomic location. Error bars are vertical 95% confidence intervals for each median pseudotime value. Late, middle, and early stages of pseudotime are indicated according to numerical y-axis values, and gene clusters are referenced. **(I**) Diagram of Bf12P1 genome showing the identified phage promoters named according to expression stage as shown in panel H. Co-expression between gene modules is indicated by the arcs colored by the calculated co-expression score.

Next, we investigated how *B. fragilis* transcriptomes changed along the infection trajectory. Differential expression analysis of the infected sample across pseudotime intervals revealed a pattern of *B. fragilis* gene expression in which certain genes were primarily expressed at early or late stages of infection (**Figure 2D- E, Supplementary** Figure 6A,C). In early pseudotime (0.2 to 0.4), we observed high expression of genes encoding RNA polymerase subunits (*rpoA*, *rpoB*, *rpoC*), a SigE sigma factor (*sigE*), the transcription termination factor NusG (*nusG*), and ribosomal proteins (*rplA* and *rplK*). Expression of these transcription- and translation-associated genes shows that host transcriptome at this stage is yet largely undisturbed and reflects the phage-induced upregulation of host cellular machinery to facilitate expression of its genome(*21*).

In the mid-pseudotime range (0.4 to 0.6), we saw an upregulation of expression of *thyB* and *dfrA* genes responsible for conversion of deoxyribonucleotide dUMP to dTMP, as well as ribonucleoside reductase (RNR) genes (*nrdJ*, *nrdD*, and *nrdG*) encoding two different classes of RNRs and representing two alternative pathways of ribonucleoside conversion to deoxyribonucleosides for DNA synthesis(*22*) (**Figure 2D-E**). Thus, in mid-infection, Bf12P1 is likely mediating conversion of NTPs into dNTPs to promote its replication. A similar gene expression pattern with strong upregulation of RNRs has been observed in a different siphoviral infection of a marine bacterium(*23*). Since Bf12P1 does not encode *nrd* genes like many other phages(*24*), it may be repurposing *B. fragilis* genes for replication of its DNA.

In late pseudotime (above 0.6), we observed only a small number of host genes that were selectively enriched, including the *priA* gene encoding a primosomal protein involved in replication of some phages(*25*). We also observed strong upregulation of transfer-messenger RNA (tmRNA) or SsrA, a component of the damaged protein degradation system(*26*, *27*)(**Figure 2D-E**). This could indicate that the phage is using SsrA for maturation of some of its proteins, or the overall damage accumulation in bacterial cells approaching lysis. Alternatively, this housekeeping RNA can be more stable than mRNA in late stages of phage infection amid widespread degradation of host transcriptome, an observation supported by literature on other phage-bacteria systems(*28*, *29*).

Observed changes in BF12P1 gene expression over pseudotime intervals corresponded well with the *B. fragilis* transcriptome changes (**Figure 2F-G, Supplementary** Figure 6B,D). Genes enriched in the pseudotime range between 0.2-0.6 were related to DNA synthesis, DNA binding, and integration, marking the period of phage replication in the host (**Figure 2F-G)**. As pseudotime values increased above 0.6, we found predominant expression of lysis, structural, and capsid maturation genes (**Figure 2F-G**).

To better understand the relationship between the expression and genomic localization of phage genes, we next calculated the median pseudotime associated with the expression of each phage gene. We then mapped the genomic co-localization of genes (**Figure 2H**). This analysis revealed that phage genes are organized in three distinct modules: early, middle, and late (**Figure 2H**, **Supplementary Table 8**). Genes within each module usually colocalized in the phage genome and were highly co-expressed (**Figure 2H,I)**. By identifying the putative phage promoters, we further resolved one gene cluster in the early gene module regulated by two putative promoters, three gene clusters in the middle module regulated by seven putative promoters, and two gene clusters in the late module regulated by five putative promoters, where gene clusters likely represent operons (**Figure 2I, Supplementary** Figures 7**, 8**).

Given the lack of known functions for the early module phage genes(16–20), we performed structural modeling with AlphaFold 3(*30*) and searched for similar protein structures, revealing that the gene 20 product is similar to the AcrIF9 anti-CRISPR protein, an inhibitor of the I-F CRISPR system(*31*) (**Supplementary Table 9**). We found that gene 20 was conserved and had homologs in 70 closely related gut phages(*32*) (**Supplementary** Figure 9). We speculate that the earliest expressed Bf12P1 genes are responsible for the initial host takeover by inhibiting its defense systems. Several conserved genes of the middle module are involved in nucleotide metabolism: gene 13 encodes putative thymidylate synthase and gene 54 encodes putative RNase E. These genes are likely involved in active conversion of host transcripts into ribonucleotide and provide substrates for host-encoded RNRs, mirroring the corresponding upregulation of these genes in the host in the middle pseudotime range. Two genes(44–45) within the late module gene cluster 2 are characterized by the latest expression in the infection trajectory (median pseudotime values >0.64) (see **Supplementary** Figure 8). These genes encode a tail protein (gene 45) and a tail terminator protein (also known as connector, gene 44), presumably involved in the latest stage of phage particle assembly(*33*, *34*).

Combined, these observations demonstrate that microSPLiT enables reconstruction of the bacteriophage infection timeline from individual cells, profiling the concurrent transcriptomic changes in the host and the phage. Moreover, the pseudotime approach reveals early and late phage gene modules, with the former responsible for the rapid disabling of the host defenses to promote infection.

## Phase-variable capsular polysaccharide (CPS) and Tsr16-adjacent region gene expression affects cellular susceptibility to bacteriophage infection

To study the effects of differences in gene expression on phage susceptibility, we examined transcriptional heterogeneity in our dataset. Within the untreated and uninfected cells, we discovered several transcriptional clusters (**Figure 3A**). A distinct cluster was marked by the expression of an operon transcribed from a phase-variable promoter activated by the tyrosine site-specific recombinase, Tsr15. This operon encodes components of type 3 fimbria that, when the promoter region downstream of Tsr15 is in the ON state, promote aggregation and biofilm formation(*35*). Another separate cluster was defined by the expression of several co-directional genes organized in an operon (BF9343_RS20580-20610). A similarity search based on predicted protein structures using the AlphaFold database indicates that the operon presumably encodes a variant of fimbriae. This operon is adjacent to genes encoding a putative AraC-family transcriptional regulator (“TR”, BF9343_RS20615) and recombinase (BF9343_RS20620, identified as Tsr16(*36*)). The TR gene is flanked by perfect 23 bp-long inverted repeats; by performing in-depth analysis of F3 sequencing data, based on reads alignment, we discovered the inversion of the TR gene in cells expressing the Tsr16-adjacent operon (**Figure 3B, Supplementary** Figure 10A). We also found that the BF9343_RS20580-20610 operon was transcriptionally active when the TR gene was co-oriented with the rest of the operon, and silent when the gene was inverted (**Figure 3B, Supplementary** Figure 10B). We conclude that the Tsr16 gene cluster represents another phase-variable region in the *B. fragilis* genome controlled by the Tsr16 recombinase.

**Fig. 3:**
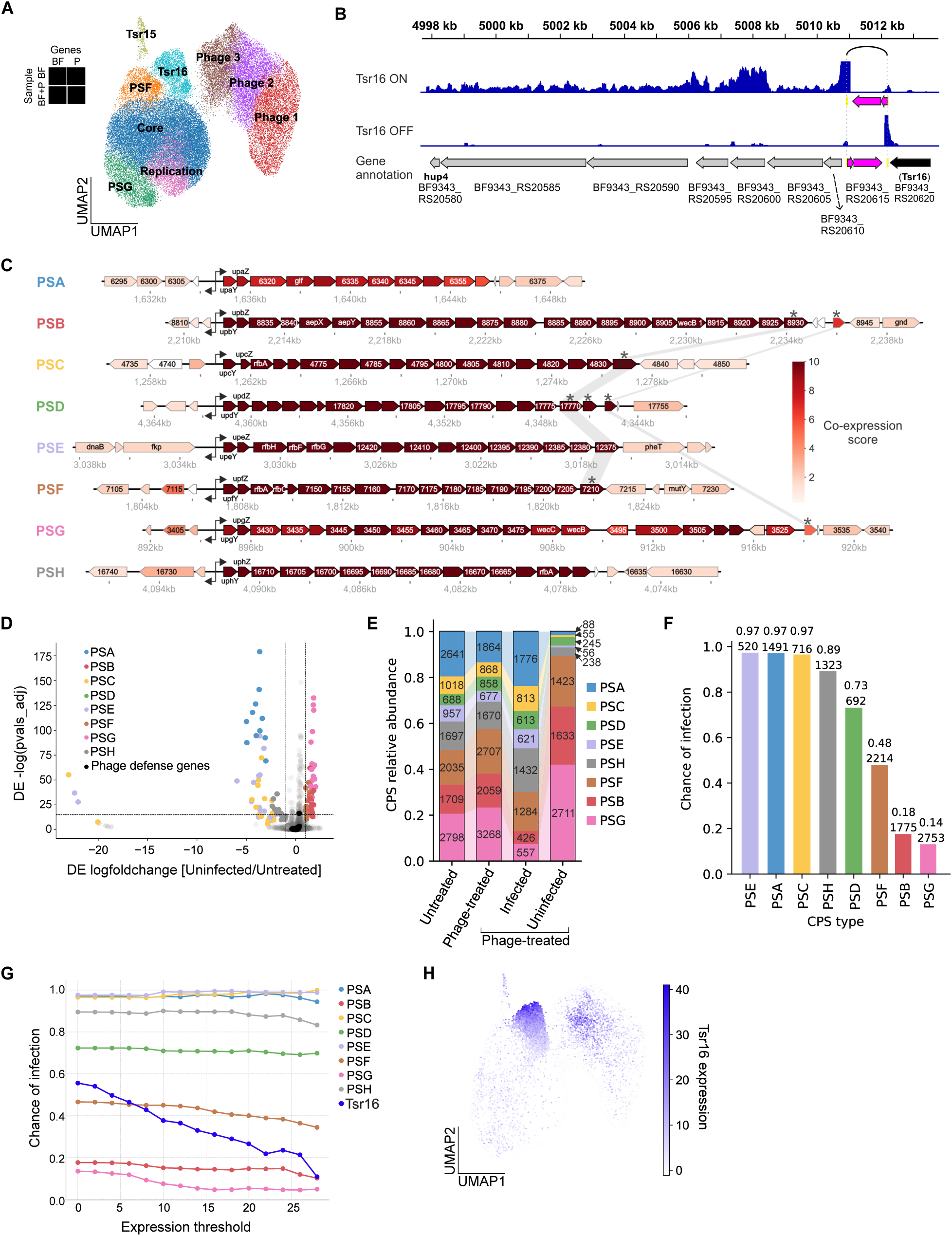
Cells expressing distinct capsular polysaccharide (CPS) types and Tsr16-adjacent region differ in chance of infection. **(A)** UMAP of grouped louvain clusters for both phage-treated and untreated samples, using both host and phage genes, labeled by gene expression patterns. **(B)** Density of microSPLiT reads for the Tsr16-associated gene cluster. Reads density is shown for the reference genome sequence (top track) and for a sequence with inverted regulatory region (bottom track). Repeat regions presumably involved in site-specific recombination are shown with pink arrows in the corresponding direction and are connected with a black arch. **(C)** Operon maps for each of 8 CPS types, with co-expression values calculated between each gene and all other genes within the operon and shown as color. Three genes flanking each operon were also included. Uniquely identified genes at the end of each operon displaying high co-expression with other genes in the corresponding operon are indicated with an asterisk. Gene paralogs with >80% sequence identity are highlighted. **(D)** Differentially expressed genes between the phage-treated, uninfected and untreated cells. Defense genes were annotated in the data using DefenseFinder(*39*). Each gene in a CPS operon is colored according to the CPS type. **(E)** Proportions of cells expressing each CPS type for untreated, phage-treated, infected, and uninfected cell subpopulations. **(F)** Infection chance of cells expressing each CPS type, calculated for each capsular variant as the proportion of infected cells in this group. **(G)** Calculated probability of infection of cells expressing either the Tsr16 gene cluster or a given CPS type above indicated expression thresholds. **(H)** Summed expression values for the Tsr16-adjacent operon shown on a UMAP embedding of all samples with both *B. fragilis* and phage transcripts.

Finally, two distinct clusters were marked by the expression of genes belonging to distinct CPS operons controlled by phase-variable promoters: namely, PSF and PSG(*9*). Overall, *B. fragilis* has eight operons encoding genes for biosynthesis of distinct capsule types, named PSA through PSH. Seven of them, excluding PSC, are expressed from a promoter which can be stochastically inverted by the Mpi recombinase and is only active in the ‘ON’ orientation. When plotting the combined expression of genes within each CPS operon on the cluster map for the entire dataset, we observed distinct patterns of expression (**Supplementary** Figure 11). We saw no co-expression between any two CPS types in the same cell (**Supplementary** Figure 12), consistent with prior literature showing that despite the frequent presence of multiple CPS promoters in the ‘ON’ state per cell simultaneously, only a single capsule type is produced in each cell(*37*, *38*). Meanwhile, genes within each operon were highly co-expressed with clear operon boundaries indicated by a drop in co-expression score (**Figure 3C, Supplementary Note 3**). The identification of multiple transcriptionally distinct subpopulations of *B. fragilis* within the untreated sample highlights the ubiquitous phase variation creating extensive phenotypic heterogeneity in this bacterium.

Since no phage load was detected in the phage-treated cells within these clusters, we hypothesized that these phase-variable regions may protect against phage infection. Capsular polysaccharide (CPS) biosynthesis genes were the most differentially expressed gene group between infected and uninfected cells, with PSF, PSB, and PSG capsular types strongly enriched in the uninfected cells (**Supplementary** Figure 13). Meanwhile, none of the known anti-phage defense systems which we annotated in *B. fragilis* using DefenseFinder(*39*), nor Tsr15 and Tsr16 regions, were differentially expressed (**Supplementary** Figures 13**, 14**).

In the comparison between uninfected and infected samples, the differentially expressed genes may reflect both the changes induced by phage and the selection of protective phenotypes from the pre-existing heterogeneous pool of states. To remove the phage-induced changes from our analysis, we compared gene expression between uninfected and untreated cells (**Figure 3D**). Again, the CPS biosynthesis genes were the strongest differentially expressed group (**Figure 3D**).

In an analysis of the distributions of cells with distinct capsular phenotypes across our dataset, we did not observe substantial differences in CPS composition between untreated and phage-treated *B. fragilis* samples, suggesting that short-term exposure to phage did not induce CPS switching. The uninfected and infected subpopulations, however, had markedly different CPS distributions (**Figure 3E, Supplementary** Figure 15), suggesting phage-mediated selection for specific capsule types. By calculating infection chances for cells expressing each capsule, we found that infection chances span a wide range between 0.14- 0.18 for the most protective capsules, PSG and PSB, to 0.97 for the most vulnerable capsules, PSC, PSA, and PSE (**Figure 3F**). Interestingly, two capsule types, PSF and PSD, had intermediate infection chances of 0.48 and 0.73, respectively, suggesting that the infection outcome is more complex than a simple selection of fully ‘vulnerable’ capsules over fully ‘protective’ ones by phage (**Figure 3F**).

We next asked whether the degree of phage susceptibility or resistance for each capsule type varied proportionally with the strength of CPS or Tsr16-adjacent operon expression. The capsule type produced, rather than CPS expression level, determined phage infection probability (**Figure 3G**). However, unlike the capsular phenotypes, Tsr16-mediated protection strongly increased in a dose-dependent manner, with infection chance dropping essentially to zero with high expression levels (**Figure 3G,H**). Fimbrial components, the predicted products of genes in this operon, could potentially act as phage decoys or promote aggregation of bacteria shielding some of them from the phage.

Finally, to determine if the discovered protective factors had a similar effect under different environmental conditions, we calculated the infection chances for cells with different capsule types from the F3 dataset samples obtained in presence of DCA. Concordant to the obtained results without DCA, we found that cells expressing PSB and PSG had low infection chances (**Supplementary** Figure 16A), and the Tsr16-adjacent operon expression protected from phage infection in a dose-dependent manner (**Supplementary** Figure 16B). Interestingly, infection chances were overall lower in presence of DCA which indicates that this bile acid may act as a protective agent, for example, affecting phage binding to the cell surface(*40*).

Taken together, these observations show that the expression of distinct phase-variable loci such as CPS operons and Tsr16 gene cluster strongly influences *B. fragilis* vulnerability to Bf12P1 infection across conditions.

## Selection for protective CPS types underlies Bf12P1 phage resistance

To experimentally confirm that certain capsule types indeed conferred protection against the Bf12P1 phage, we used *B. fragili*s NCTC 9343 ‘phase-locked’ strains unable to switch between CPS types(*41*). Specifically, we used four previously characterized strains: ‘PSA/PSE’, ‘PSB/PSG’, ‘PSD/PSH’, and ‘PSC’, referring to the CPS promoters locked in the ‘ON’ orientation in each strain(*41*). We confirmed the promoter orientations in these strains and demonstrated that on a transcriptional level each strain expresses a single dominating type of capsule: PSA in the ‘PSA/PSE’, PSG in the ‘PSB/PSG’, PSH in the ‘PSD/PSH’, and PSC in the ‘PSC’ strain (**Supplementary** Figure 17).

In a liquid culture phage infection assay, we observed that ‘PSA/PSE’, ‘PSD/PSH’, and ‘PSC’ strains, all expressing phage-sensitive capsules according to the microSPLiT data, exhibited a sharp drop in optical density and failed to recover when incubated with the Bf12P1 phage (**Figure 4A**). The wild type culture, challenged with the same phage titer, was lysed after 5-6 hours of incubation but then resumed growth, presumably due to propagation of phage-resistant bacteria (**Figure 4A**). The growth of the ‘PSB/PSG’ strain, however, was unaffected by phage. A phage plaque assay similarly showed that all tested CPS ‘phase-locked’ strains were highly sensitive to phage except for the ‘PSB/PSG’ strain, for which no plaques were observed even at the highest MOI (**Figure 4B, Supplementary** Figure 18A). These results demonstrate that expression of a capsular type predicted to be the most protective by microSPLiT data, PSG, is rendering the bacteria resistant to phage infection.

**Fig. 4:**
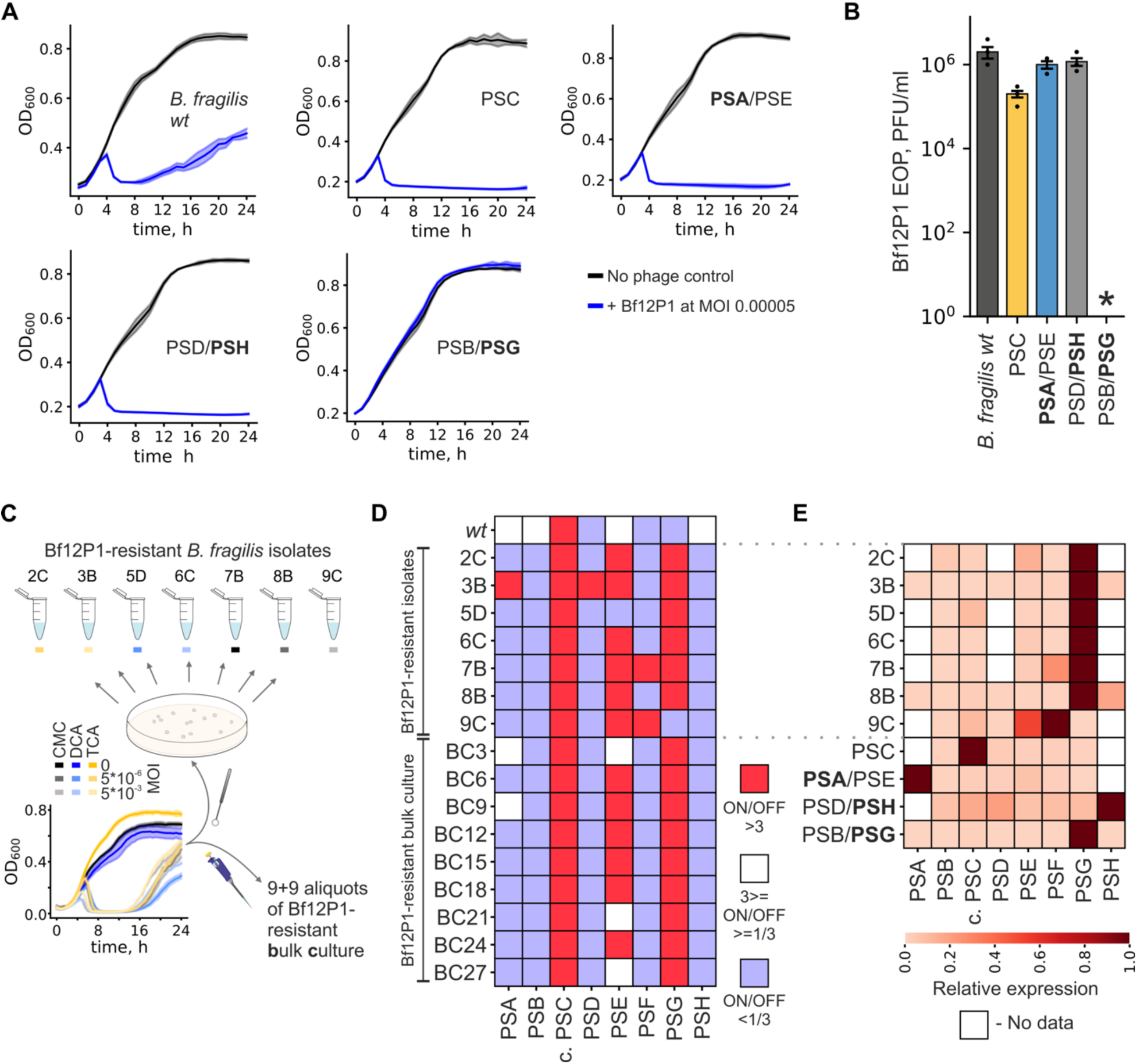
Specific capsular polysaccharide types provide resistance to Bf12P1 phage. **(A)** Growth curves for the liquid culture phage assay performed with CPS ‘phase-locked’ *B. fragilis* strains and the wild-type (*wt) B. fragilis* strain. Bf12P1 was added at time point 0 at MOI of 0.00004 (blue curves), and untreated samples are shown as controls (black curves). Shaded curves represent a 0.95 confidence interval of the mean of four biological replicates. **(B)** Quantification of a phage plaque assay performed with CPS ‘phase-locked’ *B. fragilis* strains and the *wt B. fragilis* strain. Bars, error bars, and black dots represent, respectively, the mean of three biological replicates, standard error of mean, and individual data points. Black asterisk indicates no plaques observed for the ‘PSB/PSG’ ‘phase-locked’ strain. **(C)** Schematic of an experiment for selection of Bf12P1-resistant isolates and bulk culture aliquots. **(D)** CPS promoter states in the *wt B. fragilis*, Bf12P1-resistant isolates, and bulk culture aliquots after incubation with phage at 0.005 MOI measured from WGS data. “c. PSC” denotes the non-invertible, ‘constantly ON’ promoter for the PSC operon. **(E)** Relative expression of CPS operons in Bf12P1-resistant isolates and CPS ‘phase-locked’ strains, normalized to the operon with maximal expression for every isolate or strain.

For additional confirmation, we collected phage-resistant bacteria in various conditions including the presence of bile acids (deoxycholic or taurocholic acids) (**Supplementary Note 4**, **Supplementary Table 6**). Specifically, we collected eighteen aliquots of wild-type culture (referred to later as bulk culture samples) and isolated seven colonies (referred to later as the resistant isolates) from the outgrowth following phage exposure (**Figure 4C**). Using WGS, we examined the CPS loci orientations in resistant isolates and bulk culture samples, finding that the majority of samples harbored the PSG promoter in an ‘ON’ orientation. The only exception was isolate 9C, which had the PSF promoter in the ‘ON’ state, predicted to provide ‘intermediate’ protection (**Figure 4D**, **Supplementary** Figure 18B). We confirmed that despite multiple CPS promoters in the ‘ON’ state in both resistant isolates and bulk culture samples, only PSF and PSG operons were expressed (**Figure 4E**). Since the initial culture had PSG and PSF promoters mostly in the ‘OFF’ position, these findings indicate strong selection for the protective capsular types in the presence of Bf12P1 phage.

To check whether any of the isolates acquired mutations which could be underlying resistance against Bf12P1, we performed variant calling on WGS data which revealed one non-ancestral mutation present in five resistant isolates and the majority of resistant BC samples (**Supplementary Table 6**). This mutation, however, was not present in the ‘PSB/PSG’ strain resistant to phage, suggesting it was not essential for resistance against Bf12P1. Of note, the ‘PSB/PSG’ strain did not have any additional mutations compared to the ancestral strain (**Supplementary Table 6**). We also found no reads mapped to the Bf12P1 genome in the WGS data for resistant isolates or BC samples, dismissing lysogeny with Bf12P1 as a potential resistance mechanism(*42*).

Combined, these data indicate that selection for protective capsular phenotypes, rather than accumulation of specific mutations, underlies resistance to Bf12P1.

## Combinatorial expression of anti-phage defense systems and capsules determines the phage vulnerability landscape and leads to diverse super-resistant phenotypes

Since PSF-expressing cells had ‘intermediate’ infection susceptibility compared to strongly protected PSG-expressing cells in the microSPLiT data (see **Figure 3E**), we investigated the differences in phage vulnerability between resistant isolates expressing either PSG (5D and 7B) or PSF (9C). All three isolates were resistant to lysis in liquid culture at a wide range of MOIs (**Figure 5A**). Interestingly, in a phage plaque assay, the numbers of infectious phage particles seen in isolate 7B was equivalent to the wild-type strain, despite isolate 7B expressing PSG capsule, whereas isolates 5D and 9C produced no plaques, resembling the fully protected phenotype of the ‘PSB/PSG’ strain (**Figure 5B**, **Supplementary** Figure 19). These discrepancies indicate that other mechanisms contribute to protection against the Bf12P1 phage in addition to capsular variation.

**Fig. 5:**
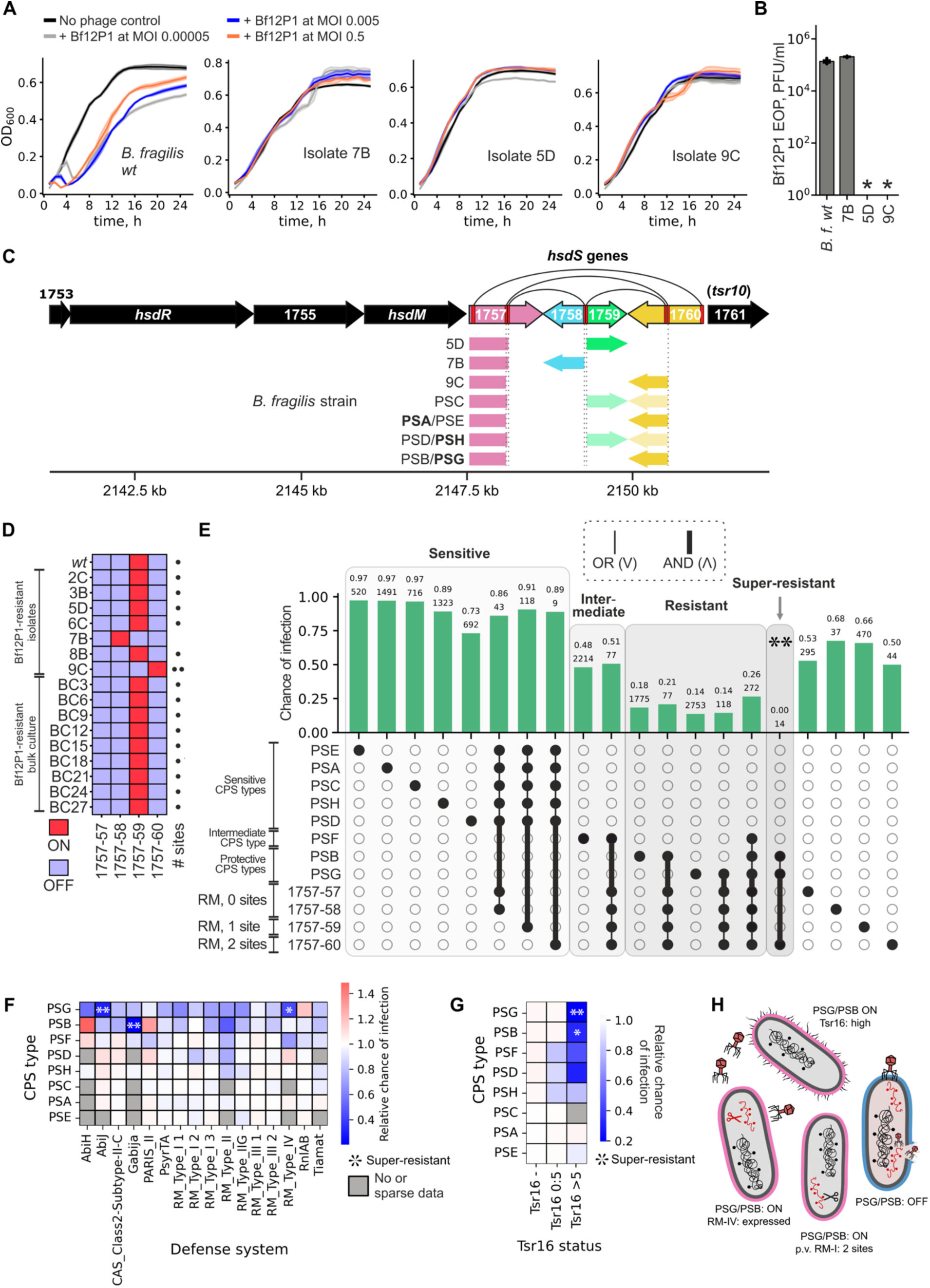
Combinatorial expression of capsule type, phase-variable regions and anti-phage defense systems creates a phage susceptibility landscape and leads to super-resistant phenotypes. **(A)** Liquid culture phage infection assay performed with Bf12P1-resistant *B. fragilis* isolates 7B, 5D, 9C, and the *wt B. fragilis* strain. Bf12P1 was added at time point 0 at indicated MOIs, black curves show untreated samples. Shaded curves represent a 0.95 confidence interval of the mean of four biological replicates. **(B)** Quantification of phage plaque assay performed with Bf12P1-resistant *B. fragilis* isolates 7B, 5D, 9C, and the *wt B. fragilis* strain. Bars, error bars, and black dots represent, respectively, the mean of three biological replicates, standard error of mean, and individual data points. Black asterisks indicate no plaques observed for the 5D and 9C isolates. **(C)** Genomic organization of the phase-variable RM type-I gene cluster. Black arches connect pairs of joint recombination sites shown with red lines. Composition of specificity subunit gene is shown below for selected Bf12P1-resistant *B. fragilis* isolates and CPS ‘phase-locked’ strains based on WGS data. Lighter colored 5’ portions of the gene for ‘PSC’ and ‘PSD/PSH’ strains indicate a mixture of states. **(D)** Phase-variable RM type-I specificity subunit states in the *wt B. fragilis*, Bf12P1-resistant isolates, and bulk culture samples (after incubation with phage at 0.005 MOI). The number of sites recognized by each version of specificity subunit in the Bf12P1 genome is shown as dots. **(E)** Phage infection chances estimated for indicated cell subpopulations based on microSPLiT data. Infection chance and number of cells in a subpopulation are shown above each bar. Sub-populations are grouped according to the level of protection against the phage. Double asterisks indicate ‘super-resistant’ cells. **(F-G)** Relative probability of infection for cells co-expressing each CPS type and **(F)** the indicated anti-phage defense mechanisms simultaneously or **(G)** Tsr16 gene cluster with indicated thresholds for expression. Subpopulations with <10 cells were excluded (labeled in gray). Combinations with double asterisks indicate ‘super-resistant’ cells (infection chance <0.05); a single asterisk represents ‘super-resistant’ cells with infection chance <0.07. **(H)** Select combinations of variable protective mechanisms leading to super-resistance to Bf12P1 infection, as opposed to vulnerable phenotypes exemplified by sensitive capsule expression.

To test whether additional phase-variable regions in the *B. fragilis* genome are involved in protection against Bf12P1, we developed an algorithm for detection of chromosomal rearrangements and applied it to WGS data for CPS ‘phase-locked’ strains, select resistant isolates, and bulk culture samples. We specifically checked whether any of the anti-phage defense systems in the *B. fragilis* genome appeared phase variable. As a result, we identified a phase-variable type-I restriction-modification (RM-I) system which is able to shuffle between 8 versions of specificity subunits by site-specific recombination(*43*, *44*) (**Figure 5C, Supplementary** Figure 20). Specificity proteins direct the RM complex to DNA recognition sites which are subsequently methylated by the complex if they are native to the host and cleaved if they are foreign substrates(*45*). Previous work identified distinct DNA recognition sequences for four of the most frequently observed specificity protein variants(*44*), revealing that the Bf12P1 genome harbored a single recognition motif for the variant 1757-59, two motifs for the 1757-60 variant, and no motifs for the 1757-57 and 1757-58 variants (**Supplementary** Figure 21A). The sparsity of occurrences of these simple 6-nucleotide motifs in the Bf12P1 genome suggests negative evolutionary pressure on these sequences (**Supplementary** Figure 21A).

Next, we investigated whether shuffling of this RM system affected vulnerability to Bf12P1. The majority of resistant isolates, including 5D, and the bulk culture samples had the 1757-59 version of a specificity subunit that recognizes a single motif in the Bf12P1 genome. Meanwhile, the 7B isolate which was sensitive to phage in the plaque assay and the resistant 9C isolate harbored the 1757-58 and 1757-60 specificity subunits for which there were no motifs and two motifs, respectively (**Figure 5D**, **Supplementary** Figure 21B). We hypothesized that the differences in these RM-I recognition sites might underlie the variation in phage susceptibility between resistant isolates.

To investigate the complementary effects of CPS and RM-I systems on a single-cell level, we calculated the infection chances for subpopulations of cells expressing distinct combinations of these systems from the microSPLiT data (**Figure 5F**). We observed that phage susceptibility remained comparable between cells of the same CPS type, regardless of whether or not they also expressed the phase-variable RM-I system in any configuration. To further analyze cells by the pattern of specificity subunit variant expression, we collected more data due to scarce detection rates of distinct specificity subunits (detected in <10 cells). For this purpose, we performed a targeted PCR-based enrichment for expression of RM-I specificity subunit variants from the same microSPLiT library. After enrichment, we resolved the expression of all four RM-I specificity variants in cells of different CPS types in our dataset, dominated by 1757-57 and 1757-59 variants. We found that the expression of these variants of specificity proteins did not additionally protect cells with sensitive capsules, resulting in infection chances above 0.7 regardless of RM-I subunit expression. However, we saw the emergence of a small group of PSB and PSG-expressing cells with the most protective specificity subunit 1757-60 that were completely uninfected (**Figure 5F**). The cells expressing this combination of traits represented a cell group we termed ‘super-resistant’, defined by a chance of infection below 0.05, the minimum expected level of non-unique cell barcoding in our microSPLiT dataset(*46*). We conclude that the protective effects of the phase-variable RM-I system emerge only in the context of a protective CPS type, which is a sign of epistasis between CPS and RM-I.

We then aimed to resolve whether the heterogeneous expression of other factors implicated in phage protection, in the context of protective CPS variants, could lead to super-resistance. Besides the phase-variable CPS and RM-I systems, *B. fragilis* NCTC 9343 encodes 15 other putative anti-phage defense systems including multiple R/M systems of types I, II, III, and IV; DNA degradation machinery (Gabija); abortive defense mechanisms (AbiJ, AbiH); toxin-antitoxin (TA) systems (PARIS, PsyrTA, RnlAB); and some systems with yet unknown mechanism (Tiamat) (**Supplementary Table 8**). Analysis of infection chances for cells of distinct CPS types expressing these systems revealed a common pattern: expression of almost all anti-phage systems slightly decreased the chance of infection when co-expressed with a protective CPS type (**Figure 5G**). Similar to the observed relationship between capsular expression and phase-variable RM-I system shuffling, this finding indicates an epistasis between CPS and other previously described defense systems.

In particular, the expression of AbiJ system simultaneously with the PSG capsule, and the expression of Gabija together with the PSB capsule, was associated with essentially no chance of infection (<0.05), thus defining cells expressing these combinations as super-resistant (**Figure 5G**). AbiJ from *B. fragilis*, classified into ‘Abi’ or abortive infection proteins, consists of N-terminal and C-terminal domains, the latter of which is a HEPN domain containing 2 catalytic residues required for nucleolytic activity(*47*). Gabija is a complex of GajA and GajB proteins involved in DNA cleavage and nucleotide hydrolysis, respectively, activated by foreign DNA binding. The cooperative interactions of these proteins allows for abortive infection defense(*48*, *49*). Additionally, cells expressing RM type-IV together with PSG had a chance of infection marginally above the threshold (0.059) and likely were also super-resistant (**Figure 5G**). The RM type-IV system contains a sole restriction enzyme that cleaves methylated DNA, in the form of foreign nucleic acids that are methylated distinctly compared to host DNA(*50*, *51*).

*B. fragilis* NCTC 9343 also encodes two putative CRISPR-Cas systems of class I subtype IB and class 2 subtype IIC, only the latter of which was expressed. WGS data of phage-resistant isolates and bulk culture samples demonstrated that a spacer perfectly matching the Bf12P1 genome was newly acquired by the CRISPR cassette associated with the IIC system in the 6C isolate (**Figure 5H, Supplementary** Figure 22). Interestingly, we found another spacer in this cassette aligning to the Bf12P1 genome with two mismatches. This partially matching spacer is present in the ancestral strain and all its derivatives, supporting the existence of a history of interactions between Bf12P1, or closely related phages, and *B. fragilis* NCTC 9343. Acquisition of a new spacer demonstrated that the CRISPR-Cas system in this strain is functional and likely provides additional adaptive immunity to bacteriophage in cells that were protected from the initial phage attack by phase variation and other defense systems.

Finally, we investigated whether other phase-variable genomic loci that were heterogeneously expressed in microSPLiT data, combined with protective capsular types, could deliver complete protection from phage. Since we detected a strong protective effect with increased expression of the Tsr16-adjacent operon (BF9343_RS20580-20610, see **Figure 3G**), we calculated the chance of infection for cells of each capsule type expressing varying levels of Tsr16 gene cluster. This analysis revealed a strong drop of infection chance in cells with high expression of Tsr16-adjacent region together with protective capsules PSG (0.034) and PSB (0.067). Though marginally above the threshold of 0.05 for the latter combination, it is reasonable to consider cells expressing both of these combinations super-resistant (**Figure 5G**).

The small size of the ‘super-resistant’ cell group (349 cells across all identified combination of traits leading to super-resistance, or 1.3% of the “Phage-treated” sample) explains the nearly complete lysis of the wild-type culture in the presence of Bf12P1, followed by slow outgrowth caused by the propagation of resistant bacteria (see **Figure 4E**). Modeling of the culture growth dynamics in the presence of phage (see **Supplementary Note 5** and **Supplementary** Figure 23) additionally supported the existence of a small number of ‘super-resistant’ cells within the culture prior to phage exposure (estimating it between 0.5-1.5% of the initial *wt* population).

We conclude that phase-variable genomic loci and heterogeneously expressed anti-phage defense systems create a pre-existing, stochastic combinatorial phage vulnerability landscape across the *B. fragilis* population, which allows a small fraction of host cells to evade phage infection without the need for acquiring mutations. Moreover, near-complete phenotypic resistance to phage (which we termed ‘super-resistance’) can be achieved by the expression of multiple combinations of variable traits, highlighting the existence of rare phenotypically discrete subpopulations of cells surviving phage exposure (**Figure 5H**). Interestingly, when we explored whether this phenotypic landscape was sensitive to environmental conditions, such as growth in a different media, we found that while the vulnerability of different CPS types to phage diminished after bile acid exposure, the shape of the overall combinatorial protective landscape remained relatively stable (**Supplementary** Figure 16).

## Discussion

Bacterial single-cell transcriptomics is a powerful tool for unraveling multiple aspects of lytic bacteriophage infection from a single sampling prior to lysis. For instance, we reconstructed the stages of bacteriophage infection using the asynchronicity of infection across single cells(*52*). Using this approach, we classified phage genes into functional modules which contained uncharacterized genes without homologs alongside genes with inferred functions. Consequently, we assigned putative function to novel phage genes, including genes that are expressed at the earliest point in infection and are presumably involved in disabling host defenses. By analyzing both the phage and host gene expression patterns concurrently as infection unfolded, we demonstrate how this approach can be used to decipher complex regulatory programs involved in host-bacteriophage interactions. Future increases in resolution of bacterial single-cell RNA sequencing may enable more insights into the intricacies of gene regulatory networks at the host-pathogen interface.

We also demonstrated the utility of microSPLiT for studying phenotypic heterogeneity in anaerobic bacterial species found in human gut microbiota, identifying, in addition to multiple phase-variable regions active in different cell subpopulations, a completely uncharacterized operon (Tsr16 gene cluster) which was heterogeneously expressed during exponential growth in culture. This operon affected phage susceptibility in a dose-dependent manner and led to super-resistance at high levels of expression in cells concurrently expressing PSG and PSB capsules. Moreover, we used microSPLiT data to predict the regulation of expression of this operon, presumably expressing fimbrial components, by an adjacent phase-variable transcription factor.

Protection from phage infection for only a few cells in a population could arise from immunity mechanisms heterogeneously induced in cells by exposure to phage. Alternatively, or in combination, it could result from pre-existing heterogeneity of cellular attributes irrespective of phage treatment, leading to different degrees of vulnerability to phage. Here, we found that the pre-existing phase-variable expression of different CPS types and either select phase-variable loci (Tsr16-adjacent operon) or anti-phage defense systems, such as RM systems type I and IV, AbiJ, and Gabija, provided near-complete protection against Bf12P1 phage. By acting together, these variable traits created a combinatorial phenotypic landscape that allowed the population to survive a phage attack by selection and propagation of a small subpopulation carrying the most protective combinations.

The novel siphovirus Bf12P1 that we isolated in this study is closely related to other recently described lytic phages targeting *B. fragilis*: VA7(*53*), vB_BfrS_23(*54*), and Barc2635(*55*) (**Supplementary** Figure 1). Of these, phage VA7 has been positively evaluated for therapeutic potential against enterotoxigenic *B. fragilis* infection in colonic epithelial cell culture models(*53*). Interestingly, mice monocolonized with *B*.

*fragilis* and Barc2635 exhibited reduced PSA capsule expression(*55*), suggesting that this related phage targets similar sensitive capsular phenotypes. As PSA is the capsule type most associated with abscess formation by *B. fragilis in vivo*(*56*, *57*), it is noteworthy that we identified PSA as the most vulnerable capsule type to Bf12P1 infection. Mapping the phage-resistant host phenotypes for diverse phages can enable the rational design of phage cocktails to prevent bacterial resistance and potentiate phage therapies.

In conclusion, our study offers a novel conceptual lens to interrogate bacterial anti-phage defense mechanisms, highlighting phenotypic heterogeneity of these systems as a primary driver of phage evasion for organisms such as gut *Bacteroidales*. The widespread phase variation throughout the *Bacteroidales* genomes ensures the existence of multiple subpopulations of cells within the same species primed to withstand attacks by multiple phages, which could be targeting cells with distinct gene expression profiles. As phage infections typically progress very quickly, this variation of a bet-hedging strategy would provide immediate protection to a subpopulation of cells expressing the most resilient phenotypes, before other, potentially slower anti-phage defense mechanisms such as CRISPR-Cas systems have an opportunity to engage. This mechanism would therefore ensure enhanced fitness for *Bacteroidales* in the unpredictable and competitive environment of the human intestinal tract.

## Methods

### Phage isolation and genome sequencing

Bacteriophage Bf12P1 was isolated from twice 0.22-µm filtered (PES membrane) primary effluent wastewater (King County Wastewater Treatment Division – West Point Treatment Plant, Seattle, WA). Phage was isolated using *Bacteroides fragilis* NCTC 9343 ATCC 25285 (hereafter referred to as *B. fragilis*). *B fragilis* was grown anaerobically at 37°C in Mega Media for 24 hours. One mL of fresh culture was added to 5mL of filtered wastewater and 5mL of fresh media. After incubating overnight at 37°C, the culture was filtered using a 0.22-µm syringe filter to remove bacterial cells. The filtrate was plated on a lawn of *B. fragilis* soft agar and incubated overnight at 37°C. The following day, potential phage isolates, which appear as zones of bacterial clearing in the soft agar (plaques), were picked and reinoculated into overnight cultures of *B. fragilis*. The following day, the cultures were re-filtered, and the filtrate was diluted and plated again onto soft agar lawns. From these plated dilutions, individual plaques, or phage isolates, were again re-amplified overnight in liquid culture. Phage isolates are stored at 4°C in glass containers in the dark. An aliquot of each potential phage isolate was used for DNA extraction and sequencing. Phage DNA was extracted using the Norgen Phage DNA Isolation kit. Phage DNA was sequenced by the Microbial Genome Sequencing Center (MiGS; Pittsburgh, PA) using Illumina NextSeq 2000. DNA sequences were quality trimmed to Q30 using the PHRED algorithm using BBDuk (BBMap version 38.92(*58*)). Reads were de novo assembled into contigs using MEGAHIT, and genome coding sequences (CDS) were predicted and annotated using Prokka v1.14.5(*59*) and Pharokka v1.7.3(*60*). Functional annotation of CDS prediction was performed manually using NCBI BLASTp v2.9.0 on the NCBI protein sequence database(*61*), using phold v0.2.0(*62*), and PhageScope(*63*). Additional annotation of unknown genes was performed by gene products modeling with AlphaFold 3(*30*) with a subsequent structural similarity search versus the PDB database(*64*) using Dali(*65*). The closest relative to Bf12P1 was determined using nucleotide BLAST search(*61*) on the nucleotide collection (nr/nt) standard database. Intergenomic similarity of Bf12P1 to its closest relative was calculated using VIRIDIC Web(*66*).

### Phage Transmission Electron Microscopy

Transmission electron microscopy was performed through the Fred Hutch Cellular Imaging Shared Resource. Bf12P1 samples were fixed in ½ strength Karnovsky’s fixative overnight at 4°C. 200 mesh Formvar/Carbon coated glow discharged grids were floated over 25 - 50ul bacterial suspension for 20 min. Grids were briefly rinsed with ½ strength Karnovsky’s, 0.1M Cacodylate buffer and 4 changes of distilled water. Grids were negatively stained with 1% uranyl acetate, wicked with damp filter paper and allowed to air dry before imaging. Samples were examined on a JEOL JEM 1400 transmission microscope (JEOL, Tokyo) at 120kV. Images were acquired with a Gatan Rio 4k x 4k digital camera system.

### Phage culture conditions and quantification

Phage Bf12P stocks were prepared by inoculating a liquid culture of *B. fragilis* with 20ul of high titer phage stock, incubating overnight to allow infection and lysis, and double 0.22um filtration of lysate. Phage stocks were immediately quantified using the soft agar overlay technique (Gratia, 1936) and stored in glass vials in the dark at 4°C. Dilutions of phage stocks were performed in sterilized spent *B. fragilis* growth media. All phage quantification for experiments was performed using the soft agar overlay method. Briefly, 7mL of mega media soft agar was combined with 500uL of an overnight *B. fragilis* culture, poured onto a hard agar plate, and allowed to solidify. Phage dilutions were spotted onto the plate (10ul). Plates were incubated for 24-48 hours, and phage were quantified and reported as plaque forming units per mL.

### Bacteria-phage growth curve (MOI infection dynamics)

For growth curve experiments used to characterize infection dynamics at varying MOIs, conditions were as follows: a 24-hour culture of *B. fragilis* was used (OD was consistently between 0.7 and 0.8). Phage stocks with known concentrations were used. Chopped meat media (anaerobe systems) was used for the following experiments: evolution of resistance; cultures used for RNA-sequencing; cultures used for single-cell RNA-sequencing. Chopped meat media was further prepared by serially filtering through 0.45uM and then 0.22uM syringe filters to remove all pieces of meat. For 96-well plate growth curves, 200ul cultures were prepared. 20uL of *B. fragilis* was added to 170uL media and 10uL of a known concentration of phage. A “Breathe-Easy’’ film was added to the top to prevent evaporation and allow gas exchange. A BioTek Epoch 2 Microplate Spectrophotometer was used to incubate at 37°C and collect OD600 measurements for 24-hours.

### Preparation of samples for single-cell RNA sequencing

A turbid 24-hour culture of *B. fragilis* was diluted in chopped meat media (anaerobe systems; filtered through 0.45 uM filter units) with Bf12P1 at an approximate MOI of 0.00004 (total PFU/total CFU as estimated by OD600) and incubated at 37°C. For bile acid samples, a final concentration of 100 uM DCA or TCA was added to the media. The collection time point was based on previous growth curves and an expected time to lysis at this MOI. Sample OD600 measurements were monitored every 30 minutes to ensure lysis had not yet occurred at the time of collection. At 2.5-3 hours and OD600 of 0.23, cells were collected on ice, centrifuged, and resuspended in 1.5mL of fixative.

### Permeabilization, polyA tailing, and reverse transcription

Fixed cells were spun down at 10,000 rcf for 10 minutes at 4C, resuspended in cold 100 mM Tris-HCl+RI (where “RI” indicates SUPERase-In RNase Inhibitor at a final concentration of 0.1 units/µM), and spun down again at the same conditions. Cells were resuspended in 250 µM permeabilization solution (0.04% Tween-20 in PBS), incubated for 3 minutes on ice, and after 1 mL addition of PBS+RI, spun down at 10,000 rcf for 10 minutes. Cells were aspirated and resuspended in 200 µM lysozyme, incubated for 15 minutes at 37C, immediately added to 1 mL PBST, and centrifuged at 10,000 rpm for 10 minutes. Samples were then resuspended in 50 µM PBS+RI per sample post-permeabilization. Cells were counted by hemocytometer and 7 million cells were taken per plate for polyadenylation. Polyadenylate polymerase I (PAP) solution was added, and each sample was incubated at 37C for 30 minutes, spun down with 1 mL PBST, and resuspended in PBS+RI. Cells were counted again for verification of the number of cells per plate.

Reverse transcription (RT) mix was created on ice with final concentrations of 1X reverse transcription buffer, 0.25 units/µM Enzymatics RNase Inhibitor, 0.25 units/µM Superase-In RNase Inhibitor, 500 µM dNTPs, 10 units/µM Maxima H Minus Reverse Transcriptase, and 7.5% PEG8000 with a final volume of 1320 µM for a 96 well plate. Reverse transcription mix was added (12 µM) to wells; cells were vortexed and 4 µM was added to PBS+RI for a final volume of 20 µM. Each sample was loaded into different wells of the reverse transcription plate to demultiplex them during analysis. Plates were incubated at 23C for 10 minutes and at 50C for 50 minutes at 500 rpm. Then, 40 µM PBST was added to each well on ice, samples were pooled in a single tube with 24.96 µM 10% Tween-20, and the total reaction was centrifuged for 10 minutes at 10,000 rpm at 4C. The sample was aspirated and resuspended in PBS+RI and disaggregated with both a 10 µm and 1 µm pluriStrainer. The composition of the 96-well plate and barcoded oligonucleotides used for RT were as previously described(*46*).

### Ligation barcoding

The ligation master mix was made on ice, containing final concentrations of 1X T4 Ligase Buffer 10X, 0.58 units/µM Enzymatics RNase Inhibitor, 0.05 units/µM SUPERase-In RNase Inhibitor, 7.5% PEG8000, and 8 units/µM T4 DNA Ligase, with a final volume of 2040 µM. Cells were added to the ligation mix and 40 µM were transferred into each well of a Round 2 ligation barcode plate before incubating for 30 minutes at 37C and adding 10 µM of round 2 blocking solution (26.4 µM barcode 0340, 2.5X Ligase Buffer). The plate was incubated again for 30 minutes at 37C, and cells were pooled, disaggregated, and barcoded again (round 3 blocking solution: 11.5 µM barcode 0066, 125 mM EDTA). After the final filtering step, cells were washed (4000 µM 1X PBS, 40 µM 10% Tween-20, 10 µM Superase-In RNase Inhibitor) and aliquoted into libraries with 1X PBS+RI to a final volume of 50 µM. Again, the compositions of the 96-well plates and oligonucleotides used for barcoding step were as previously described(*46*).

### Sequencing library preparation

Aliquoted libraries were mixed with 50 µM lysis buffer (20 mM Tris pH 8, 400 mM NaCl, 100 mM EDTA pH 8, 4.4% SDS) and 10 µM of 20 mg/mL Proteinase K for protein digestion prior to incubation at 55C for 2 hours with 500 rpm shaking. The resulting cDNA was purified using MyOne C1 Dynabeads. For template switching, a common 5’ sequence was added to full-length cDNA by resuspending purified product in 200 µM Template Switch Mix (5X Maxima RT Buffer, 50% PEG8000, 10 mM of each dNTP, Superase-IN, 100 µM TSO, Maxima RT RNase H minus enzyme) and incubating at room temperature for 20 minutes at 600 rpm before further incubation at 42C for 90 minutes with agitation. Then, samples were washed in 250 µM Tris-T + RI, decanted and resuspended again with 100 µM per sample of terminal deoxynucleotidyl transferase tailing (TdT) master mix (10X TdT buffer, CoCl2, 10 mM dCTP, 1 mM ddCTP, 20 units/µM TdT enzyme, 5 units/µM RNase H). Following incubation for 1 hour at 37C and a wash with Tris-T + RI, cDNA was amplified via qPCR using primers BC0108 and M0002, fragmented, and ligated with adapters prior to Illumina Sequencing as was described previously(*46*).

### Data preprocessing

Sequencing was performed on NovaSeq X Plus Series (PE150, Novogene), and a genome was generated with STAR(*67*) using reference genomes for both *Bacteroides fragilis* NCTC 9343 and BF12P1. With the generated genomes, barcode lists for each of three indexing rounds, and output sequencing files, STARsolo(*68*) was run for genome alignment, preserving only the uniquely aligned reads. The raw count matrix was loaded into AnnData(*69*); after assigning reads to barcode combinations and sample wells, log-log barcode rank plots were used to determine an initial filtering threshold of 400 UMIs/cell for a reduced likelihood of doubly labeled cells. Remaining ribosomal (r)RNA and transfer (t)RNA reads were filtered out, and the dataset was filtered again by 40 UMIs/cell.

### Single-cell analysis

Two separate libraries were concatenated after filtering, log-normalized by 600 counts per cell, and scaled to unit variance and zero mean. Subsequently, ComBat(*70*) was run for batch effect correction. Normalized expression data was dimensionally reduced using principal component analysis (PCA). Shared neighbor graphs and uniform manifold approximation representations (UMAP(*71*)) were calculated with the first 12 principal components. All subsequent calculations were run in Python using Scanpy(*72*) documentation for single-cell analysis.

### Differential gene expression analysis

Scanpy gene ranking functions (sc.tl.rank_genes_groups and sc.get.rank_genes_groups_df) were used to analyze and retrieve statistical data between two groups of interest within the annotated data object. The output parameters included names of all genes, z-score, log fold change, p-values, and adjusted p-values. To create a volcano plot from this data, either the |score| or -log(adjusted p-value) was plotted on the y-axis against the log-fold change on the x axis. Lists of defense genes from DefenseFinder(*39*), host response genes upregulated after phage treatment, and CPS genes were used to assign colors to each point. Host response genes were taken as the top 15 distinct genes identified in the phage-treated sample only clustering analysis relative to the untreated sample. Initial CPS gene annotations were mapped by and received from Dr. Laurie Comstock (University of Chicago).

### Calculation of co-expression scores

To assess the co-expression between two genes A and B, first, probabilities of expression of individual genes were calculated as fractions of cells having above-zero normalized non-scaled expression values - p(A) and p(B), respectively. Then, the probability of simultaneous expression of the two genes, p(A&B), was calculated as a fraction of cells having above-zero normalized non-scaled expression values for both inspected genes. Co-expression score was calculated as a ratio of p(A&B) to the multiplication of p(A) and p(B). Values close to 1 indicate independent expression of genes; values above 1 indicate co-expression, and values below 1 indicate mutually exclusive expression of genes. To define a border of a CPS operon, mean co-expression values were calculated between genes adjacent to the CPS operon and core genes of a CPS operon. Co-expression between gene sets (e.g., CPS operons) was calculated similarly to genes, with probability of expression calculated as a fraction of cells having above-zero normalized non-scaled expression values for any of genes within a set.

### Diffusion pseudotime analysis

The data was first subsetted into the phage-treated sample alone. A root cell was defined using adata.uns[‘iroot’] = np.flatnonzero(adata.obs[‘louvan’]==0)[1], which selected a random indexed cell from the untreated cluster within all phage-treated cells. Scanpy diffusion maps were created prior to running the existing diffusion pseudotime tool with 0 branchings and 10 diffusion components. For downstream analysis using pseudotime values, 5 bins of equal size were created to group cells into pseudotime ranges (0.0-0.2, 0.2-0.4, 0.4-0.6, 0.6-0.8, and 0.8-1.0). Differential expression analysis was run to identify the top 10 distinct *B. fragilis* genes within each pseudotime bin, and the Clusters of Orthologous Groups (COGs) database was used to define functional categories for these genes. Duplicate genes were removed. Raw mean expression for each of these genes, grouped by functional category, was calculated using the adata.raw.X matrix.

### Verification of capsular operons promoter state via PCR

The procedure was adapted from the Troy et al. study(*73*) characterizing ON and OFF promoter states for the PSA locus, in addition to the recommended PCR protocol for use of KAPA Biosystems HiFi HotStart Ready Mix containing a type B DNA polymerase, optimized buffer, MgCl2, and dNTPs. In the anaerobic chamber, *B. fragilis* glycerol stocks were streaked onto blood agar plates and incubated at 37°C overnight. A single colony was transferred to 1 mL of BHI broth and grown overnight at 37°C in the chamber; 100 uL of culture was then transferred to 10 mL of BHI broth into a sealed Balch-type tube and grown to an OD600 of 0.5-0.6 at 225 rpm, 37°C. After growth, DNA was extracted from each sample using the DNA extraction protocol provided by Qiagen and concentration measured via Qubit v4.

For PCR, all forward and reverse primers were used based on Troy et al. study(*73*). Primers were resuspended to 100 uM. The PCR mix contained 12.5 uL 2X HotStart Ready Mix from KAPA Biosystems, 0.75 uL of 10 uM forward primer, 0.75 uL of 10 uM reverse primer, 1.0 uL template DNA, and remaining molecular biology grade water to a final volume of 25 uL. For all CPS promoters, annealing temperature was 55-59°C and set at 60°C; for PSD, annealing was at 67°C and set to 70°C. The protocol was set to 95°C denaturation (3 minutes), 98°C denaturation (20 sec), the respective annealing temperature (70 sec), 30X cycles, and a 15-minute extension at 72°C. The Monarch PCR cleanup kit was then used for purification and 1.0 ng of each DNA product was visualized by TapeStation.

Purified PCR products were digested at a concentration of 250 ng with 0.5 uL of the respective restriction enzyme (PSA: SspI-HF; PSB: RsaI; PSD: DraI; PSE: SspI-HF; PSF: EarI; PSG: PacI; PSH: SfcI)(*73*), 2.5 uL digest buffer, and molecular biology grade water to a final volume of 25 uL for 2 hours at 37°C. Post-digestion, products were mixed with gel loading dye and run on a 1.5% agarose gel at 100V for 30 minutes with 1 Kb standard DNA ladder. Bands were visualized and quantified on the gel to verify CPS promoter orientation for each of the phase-locked *B. fragilis* mutants.

### RM-I transcripts enrichment and data analysis

RM-I transcripts were enriched by PCR using a forward primer specific for BF9343_RS21605 (BF9343_1757) transcript and a universal reverse primer for a barcode adapter (**Supplementary Table 1**). For the first qPCR, 1.6 uL of 10 uM forward and reverse primers were used, in addition to 25 uL 2X Kapa HiFi Hotstart Ready Mix, 2.5 uL 20X evagreen, 1.0 uL of the F3 12K TdT cDNA template library, and 17.8 uL molecular biology grade water for a total reaction volume of 50 uL. PCR conditions were as follows: 3:00 at 98C hold, 33X cycles of 0:12 at 98C denaturation, 0:15 at 55C annealing, and 0:10 at 72C elongation. Then, 0.6-0.8x double-sided size selection was performed using SPRI beads; 45 uL product from qPCR reaction 1 was mixed with 27 uL SPRISelect beads, incubated for 5 minutes at RT, and separated using a magnetic stand. The supernatant (65.7 uL) was transferred to a new tube containing 9 uL beads, pipetted 10x, and incubated again for 5 minutes at RT. Supernatant was discarded. Subsequently, 180 uL of 85% fresh EtOH was added, incubated for 30 seconds at RT with the sample, and then discarded carefully. Beads were air-dried for 30 seconds to 1 minute until visibly dry. Finally, 50.5 uL molecular biology grade water was added, mixed well, and incubated for 10 minutes at RT. Size-selected samples were transferred to a new tube. Following purification, 18.5 uL of the resulting library was put into a qPCR 2 mix containing 25 uL 2X Kapa Hifi Master Mix, 2.0 uL each of our 10 uM indexing barcodes 27 and 78 (the latter for identification of the library within a mix of libraries sent for sequencing), and 2.5 uL of 20X evagreen. The PCR conditions were set as follows: 95C for 3:00 hold, 4-6X cycles of 98C for 0:20, 67C for 0:20, and 72C for 3:00, and a final hold at 72C for 5:00. After another 0.6-0.8x size selection, the final library concentration was measured, and the library sent for Illumina sequencing. Sequencing was performed in a PE 150+150 mode using Illumina NovaSeq X Plus Series at Novogene.

Obtained forward reads were aligned to a synthetic reference sequence prepared by concatenation of four variants (1757-57, 1757-58, 1757-59, and 1757-60) of a specificity subunit gene region that contains recombination sites and is involved in shuffling. For unambiguously aligned reads which overlap the repeat regions, barcode information was extracted from corresponding reverse reads. Barcode combinations found in at least 3 reads were used to additionally assign RM specificity subunits to cells in the original microSPLiT data. In case of simultaneous detection of several different specificity subunits for a barcode combination, an epistatic model was considered: 1757-60>1757-59>1757-57, 1757-58.

### Verification of capsule expression via RT-qPCR

Two pairs of forward and reverse PCR primers were designed in SnapGene to target downstream regions of each CPS operon (**Supplementary Table 1**). All primers and products were verified using *B. fragilis* genomic DNA prior to experiments with phage-resistant isolates and phase-locked mutant strains. All qPCR runs were performed with RNA controls for each of the strains as a relative comparison to cDNA samples, as they may have later amplification cycles from gDNA contamination.

Overnight cultures with 4 mL basal metabolic broth were started for each of the samples, directly from original glycerol stocks, using wild-type *B. fragilis* as a control. Cultures were incubated at 225 rpm and 37°C overnight in Balch-type tubes to an approximate OD600 between 0.6-1.0. Approximately 1.5-2.0 mL of culture was spun down at 10,000 rpm for 30 seconds and supernatant was discarded. A general TRIzol protocol from Invitrogen was followed to extract RNA from each culture pellet. Briefly, 1 mL of TRIzol was added to approximately 10^6^ cells for each sample, homogenized, and incubated for 5 minutes on ice for complete dissociation of the nucleoproteins complex. Next, 0.2 mL of chloroform was added, mixed via shaking, and incubated for an additional 3-4 minutes on ice. Samples were centrifuged for 15 minutes at 12,000xg and the aqueous phase containing RNA was transferred to a new tube. To precipitate the RNA, 0.5 mL of isopropanol was added to the aqueous phase and incubated for 10 minutes on ice. Samples were centrifuged for 10 minutes at 12,000xg, then kept at -20°C for 30 minutes and spun down again at the max rcf (approximately 21,000xg) for 20 minutes. Supernatant was discarded carefully via pipet and the pellet was resuspended in 1 mL of 75% EtOH by vortex. Then, samples were centrifuged for 5 minutes at 7,500xg and the pellets were air dried for 5-10 minutes. RNA was finally solubilized via resuspension in 25 uL RNase-free water and incubated at 55°C for 15 minutes before Qubit concentration measurements. All samples were treated with 2 uL of DNase I from New England Biolabs to a final volume of 100 uL and purified using the NEB Monarch RNA cleanup kit with an elution volume of 30 uL.

For first-strand cDNA synthesis, the standard protocol from ThermoFisher Scientific using Maxima H Minus Reverse Transcriptase was used. The total reaction volume was tripled to include 20 uL template RNA, 3 uL of 100 uM random hexamer primer stock, 3 uL of 10 mM dNTP mix, 12 uL 5X RT buffer, 1.5 uL Superase RNase inhibitor, 1 uL Maxima Reverse Transcriptase, and nuclease-free water to a final volume of 60 uL. Reaction mixture was incubated for 10 minutes at 25°C, 30 minutes at 50°C, and terminated for 5 minutes at 85°C. For qPCR, 2 replicates were run for each cDNA sample along with 2 replicate RNA samples and non-template controls. The qPCR program was set to 98°C (3 min, then 12 sec), 65°C (15 sec), 72°C (10 sec), and 33 cycles. For analysis, the number of amplification cycles for each B. fragilis sample and every CPS primer pair were taken directly from qPCR data.

### Variant calling

Variant calling was performed using snippy v 5.2(*74*) with default conditions using the *Bacteroides fragilis* NCTC 9343 GCF_000025985.1 ASM2598v1 as a reference genome. Medium-quality variants having SNP score>100 were extracted from the raw.vcf data.

### Identification of CRISPR spacer acquisition from WGS data for B. fragilis isolates

WGS reads were *de novo* assembled using SPAdes v3.15.5(*75*) with default conditions. CRISPR cassettes were predicted in the obtained contigs using Minced v0.4.2(*76*, *77*).

### Evolution of phage-resistant isolates

*B. fragilis* seed cultures were grown and prepared for the experiment as previously described. To prepare the experimental culture conditions: 20uL of overnight *B. fragilis* culture was added to 170uL twice-filtered chopped meat media and 10uL of phage to achieve final MOIs of 0.000005 (“Low-MOI”) and 0.005 (“High-MOI). Additionally, two different bile acid growth conditions were generated by adding taurocholic acid (TCA) or deoxycholic acid (DCA) or a solvent-control (methanol) to cultures at a final concentration of 100uM. “Breathe-Easy” film was added to the top of the flat-bottom 96-well plate to prevent evaporation and allow gas exchange. A BioTek Epoch 2 Microplate Spectrophotometer was used to incubate at 37°C and collect OD600 measurements for 24-hours. After 24 hours, evolution of resistance was visually confirmed by the re-growth of cultures treated with phase after the lysis/dead stage. Aliquots of bulk culture were collected via centrifugation and stored at -20C for shotgun sequencing. To isolate individual resistant isolates, an aliquot of each 24-hour culture was struck out on solid agar Mega Media plates and incubated at 37°C for 48 hours. After 48-hours, 4 individual colonies from each condition of interest were picked using an inoculation loop and inoculated into Mega Media and incubated at 37°C for 48 hours. After 48 hours, bacterial cultures were tested for resistance against Bf12P1 using a cross-streaking assay. Briefly, 20ul of high-titer phage stock Bf12P1 was dispensed at the top of an agar plate in triplicate. The plate was tilted at a 45-degree angle, allowing the aliquot to run down the plate. Phage streaks were then allowed to dry. Plates were turned perpendicular and 12ul of bacterial culture was applied in the same manner; effectively, cross streaking with phage. Plates were then incubated at 37°C for 48 hours. Isolates were determined to be resistant or sensitive by a visual examination of if they grew across the phage streak. Resistant and sensitive isolates were further confirmed to be resistant through liquid growth curve assays with Bf12P1as previously described. Glycerol stocks were prepared from individual colonies, which were also used to inoculate liquid cultures to prepare for whole genome sequencing. Turbid cultures were pelleted via centrifugation, supernatant removed, and DNA extracted. Samples were sequenced by the Microbial Genome Sequencing Center using their Illumina Whole Genome Sequencing service.

## Data and code availability

All raw sequencing data will be submitted to the SRA (Sequence Read Archive), while the processed data will be submitted to GEO (Gene Expression Omnibus) with the accession numbers provided upon publication. Code will be published and made openly available on GitHub: https://github.com/anika-gupta-8/B_fragilis_phage_microSPLiT upon publication.

## Acknowledgements

A.K. and G.S. acknowledge support from the Department of Energy Office of Science, Biological and Environmental Research (BER) Program, Grant DE-SC0023091. A.K. is supported by the National Institute of Dental and Craniofacial Research of the National Institutes of Health Grant R21DE032890, and by the National Institute of General Medical Sciences of the NIH Grant R35GM150994. N.M. was supported by a Washington Research Foundation Postdoctoral Fellowship. This research was supported by the Electron Microscopy Shared Resource RRID:SCR_022611 of the Cancer Consortium of Fred Hutch, University of Washington, and Seattle Children’s Hospital (P30 CA015704). We thank Steve MacFarlane for assistance with electron microscopy. We thank Jason Saba and Robert Landick (University of Wisconsin-Madison) for providing CPS ‘phase-locked’ strains.

## Author contributions

A.G., N.M., D.S., G.S., N.D., and A.K. designed experiments; A.G., N.M., D.S., N.L., K.G., A.R.

performed experiments; A.G., N.M., D.S., and A.K. analyzed data; D.S. and Y.I. performed mathematical modeling, G.S., N.D. and A.K. provided supervisory support; A.G., N.M., D.S., G.S., N.D., and A.K. wrote the manuscript.

