## Supplementary Notes & Figures for "Combinatorial phenotypic landscape enables bacterial resistance to phage infection"

**Affiliations:**
[1]Institute for Systems Biology, Seattle, WA, USA.
[2]Molecular Engineering and Sciences Institute, University of Washington, Seattle, WA, USA.
[3]Translational Science and Therapeutics Division, Fred Hutchinson Cancer Center, Seattle, WA, USA.
[4]Molecular and Cellular Biology Graduate Program, University of Washington, Seattle, WA, USA.
[5]Department of Physics, Center for Interdisciplinary Research in Astrophysics and Space Science, University of Santiago, Chile.
[6]Department of Electrical and Computer Engineering, University of Washington, Seattle, WA, USA.
[7]Paul G. Allen School of Computer Science and Engineering, University of Washington, Seattle, WA, USA.
[8]Microbiome Research Initiative, Fred Hutchinson Cancer Center, Seattle, WA, USA.
[9]Department of Medicine, Division of Gastroenterology, School of Medicine, University of Washington, Seattle, WA, USA.

†These authors contributed equally to this work.

### Supplementary Text

#### Supplementary Note 1. Initial analysis of *B. fragilis* + Bf12P1 sequencing data

We loaded each of the phage-treated and untreated samples into a separate set of wells in the first barcoding plate to use the first barcode as a sample identifier. Our first experiment, named "M30", yielded a total of 15,260 cells and recovered medians of 26 and 8 *B. fragilis* mRNA transcripts per cell, as well as 0 and 23 phage transcripts per cell, for the untreated and phage-treated samples, respectively (**Supplementary Figure 2A-B**). The 'barnyard' plot where we aligned all cells to both *B. fragilis* and *B. subtilis* genomes showed no cells aligning to both genomes which would represent doublets (**Supplementary Figure 2D**). Upon data examination and clustering by gene expression, we observed clear separation of phage-treated and untreated samples, as well as the segregation of *B. fragilis* into phenotypically heterogeneous clusters (**Supplementary Figure 2C**). However, the *B. fragilis* transcript abundance measured in this experiment was relatively low (**Supplementary Figure 3B**).

Next, in order to maximize the retrieval of transcripts, we implemented several modifications to microSPLiT, detailed in the Materials and Methods. Briefly, we loaded more bacterial cells (10 million) into the reverse transcription plate to avoid excessive unused primer carry-over, controlled for lysozyme unit activity, and added a TdT-based poly-dC tailing step during library preparation to append the 3' adapter to the cDNA that previously failed to acquire one through template switching. We performed library preparation on two separate subsets of barcoded cells from this experiment, which showed similar gene expression patterns and were subsequently merged into a single dataset (**Supplementary Figure 3A**). After filtering out low quality cells, this dataset ("F3") contained 26,649 and 26,643 cells from the untreated and phage-treated samples, respectively (**Figure 1I**). Since we retrieved substantially more cells with high transcript content from the F3 experiment (**Supplementary Figure 3B**), from here on we proceeded with the analysis of F3 data.

#### Supplementary Note 2. "Responders" cluster gene expression

The "responders" cluster (shown in **Figure 1L**) was marked by expression of genes of several putative ribonucleoside reductases (*nrdD*, *nrdG*, and *nrdJ*) in addition to carbohydrate metabolism genes (BF9343_RS14805, *fkp*, and BF9343_RS00795) and the *priA* gene encoding the primosomal protein involved in the restart of stalled replication forks. Further, BF9343_RS17000-BF9343_RS17005 operon was upregulated, encoding, respectively, putative thymidylate synthase (*thyB*) and dihydrofolate reductase (*dfrA*). Other uniquely upregulated genes included BF9343_RS20235, an aminopeptidase P family protein, an NADP dependent oxidoreductase (BF9343_RS04005), and proteins involved in membrane transport (BF9343_RS20250 and BF9343_RS16815, which likely is a part of the operon also containing *nrdD* and *nrdG* genes with locus tags of BF9343_RS16805 and BF9343_RS16810, respectively).

#### Supplementary Note 3. Identification of additional genes in the CPS operons based on co-expression analysis

We identified four additional genes at the ends of the previously defined PSB (BF9343_RS08930 gene), PSC (BF9343_RS04835 gene), PSD (BF9343_RS17770 gene), and PSF

(BF9343_RS07210 gene) operons that were highly and uniquely co-expressed with the rest of the operon genes. Notably, these genes were paralogous to the BF9343_RS12375 gene located at the end of the PSE operon, and encode family 4 putative glycosyltransferases that share >79% sequence identity. This further supports their likely involvement in capsular polysaccharide biosynthesis. Further, we found the BF9343_RS21660 gene (encodes an N-acetylmuramidase domain-containing protein) at the end of the PSD operon which was highly co-expressed with the rest of the operon genes. We also identified another group of three homologous genes at the ends of PSB (BF9343_RS08940 gene), PSD (BF9343_RS17760 gene), and PSG (BF9343_RS03530 gene) operons that were co-expressed with the rest of the operon genes. These genes encoded putative HU family DNA-binding proteins (share >74% of identity) which are potentially involved in regulation of CPS operon transcription. Altogether, by using the co-expression analysis, we extended CPS operons by eight genes (**Figure 3C** asterisks).

#### Supplementary Note 4. Selection of phage-resistant isolates of *B. fragilis*

Selection for phage-resistant bacteria was conducted at two different MOIs and in the presence of two bile acids (deoxycholic or taurocholic acids) which were added to investigate resistance determinants under varying media conditions. Bulk culture samples and resistant isolates were obtained from outgrowth after a 24-hour incubation with phage. Additionally, we found a random colony isolated in no-phage conditions but showed resistance (isolate 7B). Isolation conditions for bulk culture samples and resistant isolates can be found in **Supplementary Table 6**.

#### Supplementary Note 5. Modeling of *B. fragilis* culture growth with phage

To estimate parameters of phage infection, we first assumed that the initial concentration of resource is sufficient to not constrain the growth of bacteria until lysis (**Supplementary Figure 23A**). It allowed us to ignore the resource dynamics and to simplify the description of bacterial growth. We also assumed that the population dynamics of bacteria can be approximated as consisting of two phases: first, a "phage-free" growth with a constant birth rate, and subsequently, lysis when the concentration of phage becomes sufficiently high. The latter approximation will later be shown to be self-consistent and justified by the extremely rapid double-exponential phage replication. During phage-free and resource-unlimited growth, the dynamics of bacterial density *B(t)* is given by

$$\frac{dB}{dt} = \beta B\,, \quad (1)$$

$$B = B_0 e^{\beta t}\,,$$

where β is the bacterial growth rate constant determined from the initial part of the phage-free growth curve (**Supplementary Figures 23C-D**).

During the lysis phase, bacterial dynamics include the negative term describing attacks by phages with the concentration *P(t)*. This term is proportional to the bacterial replication rate because only actively dividing bacteria are susceptible to phages. The proportionality constant δ characterizes the attack rate.

$$\frac{dB}{dt} = \beta B(1 - \delta P). \quad (2)$$

The lysis time t* is defined as the moment when the bacterial growth rate becomes negative:

$$P(t^*) = \frac{1}{\delta}. \quad (3)$$

The phage growth rate is proportional to the attack term in eq. 2, with the proportionality constant α dependent on the phage burst size and latency time:

$$\frac{dP}{dt} = \beta B\delta P\alpha, \quad (4)$$

$$P(t) = P_0 e^{\beta\delta\alpha \int_0^t B(t')dt'},$$

where

$$\int_0^t B(t')dt' = B_0 \int_0^t e^{\beta t'}dt' = \frac{B_0}{\beta}(e^{\beta t} - 1) \approx \frac{B(t)}{\beta}. \quad (5)$$

Hence,

$$P(t) \approx P_0 e^{\delta\alpha B(t)}. \quad (6)$$

At the time of lysis

$$P_0 e^{\delta\alpha B(t^*)} = \frac{1}{\delta}, \quad (7)$$

$$ln(\delta P_0) + \delta\alpha B(t^*) = 0,$$

$$B(t^*) = \frac{-ln(\delta P_0)}{\delta\alpha}.$$

This explains the linear dependence of bacterial optical density at the time of lysis on the logarithm of the initial concentration of phages (**Supplementary Figure 23B**) (see also (*1*)). The slope and intercept of the regression line of the semi-log plot (**Supplementary Figure 23B**) allowed us to determine the constants δ (phage attack rate) and α (a combination of phage burst size and latency time).

Next, using estimated parameters, we modeled a long-term dynamic of the *wt B. fragilis* population that consists of a mixture of phage-sensitive $B_{Sus}(t)$ and phage 'super-resistant' $B_{Fp}(t)$ cells in the presence of phage *P(t)* and a limited resource *N(t)* as follows:

$$\frac{dN}{dt} = -\beta_{Sus}B_{Sus}(t)N(t) - \beta_{Fp}B_{Fp}(t)N(t), \quad (7)$$

$$\frac{dB_{Sus}}{dt} = \beta_{Sus}B_{Sus}(t)N(t) - \beta_{Sus}B_{Sus}(t)N(t)\delta_{Sus}P(t), \quad (8)$$

$$\frac{dB_{Fp}}{dt} = \beta_{Fp}B_{Fp}(t)N(t) - \beta_{Fp}B_{Fp}(t)N(t)\delta_{Fp}P(t) \ , \qquad (9)$$

$$\frac{P(t)}{dt} = \beta_{Sus}B_{Sus}(t)N(t)\delta_{Sus}\alpha_{Sus}P(t) + \beta_{Fp}B_{Fp}(t)N(t)\delta_{Fp}\alpha_{Fp}P(t) \ , \qquad (10)$$

Eq. 7 describes the consumption of non-renewable resource *N(t)* by phage-sensitive and super-resistant bacteria.

We estimated the growth rate constants $\beta_{Sus}$ as the averages for the phage-sensitive CPS 'phase-locked' strains (PSC, PSD/PSH, PSA/PSE) and for *wt B. fragilis* (**Supplementary Figure 23C**). For the 'super-resistant' population, we used the average of growth rate constants $\beta_{Fp}$ estimated for the PSB/PSG CPS 'phase-locked' strain and for phage-resistant isolates (**Supplementary Figure 23D**).

Eq. 8-9 describe the population dynamics of susceptible (phage-sensitive) and 'super-resistant' cells. Since we chose to measure the resource by the number of cells it can support, the growth terms in eq. 8-9 are identical to the loss terms in eq. 7. Since it is assumed that phages attack only actively dividing cells, the death terms in eq. 8-9 consist of the cell birth rate multiplied by the phage concentration and the attack rates $\delta_{Sus}$ and $\delta_{Fp}$.

Eq. 10 describes the dynamics of the phage population. The phage population is assumed to grow monotonously with a rate equal to that of bacterial death multiplied by the factor α that, as in eq. 4, accounts for the phage burst size and latency time. The phage constants $\delta_{Sus}$ and $\alpha_{Sus}$ for the phage-sensitive population were obtained as described above using the data for *wt B. fragilis* (**Supplementary Figure 23B**). For 'super-resistant' cells, $\delta_{Fp}$ was set to 0 to reflect absolute protection of these cells against the phage. Values and descriptions of model parameters are summarized in **Table 1**.

**Table 1**. Values and descriptions of model parameters that were used to solve the system of differential equations. The analysis (eq. 1-7) is based on the approximation that phages do not affect bacterial growth until sudden lysis, so the parameters obtained using the fit given by Eq. 7 are approximate as well. To improve the accuracy of the model, the values of α were adjusted to better reproduce the timing of the maximum bacterial population and the beginning of massive lysis. This explains #multiplication by factor 2 shown in the table.

| Parameter | Value | Description |
|---|---|---|
| $\beta_{Sus}$ | 2.99e-11+/-4.0e-12 (STD) mL/h*CFU | Bacterial growth rate constant for phage-sensitive cells |
| $\beta_{Fp}$ | 2.56e-11+/-3.8e-12 (STD) mL/h*CFU | Bacterial growth rate constant for phage 'super-resistant' cells |
| $\delta_{Sus}$ | 3.12e-11 mL/PFU | Phage attack rate for phage-sensitive cells |
| $\delta_{Fp}$ | 0 mL/PFU | Phage attack rate for phage 'super-resistant' cells |
| $\alpha_{Sus}$ | 61*2# PFU/CFU | Combination of phage burst size and latency time for phage-sensitive cells. |
| $\alpha_{Fp}$ | 61*2# PFU/CFU | Combination of phage burst size and latency time for phage 'super-resistant' cells |

Interestingly, the growth rate estimated for phage-resistant *B. fragilis* strains and isolates was 14% lower than the growth rate for phage-sensitive strains, indicating an additional fitness burden of phage resistance.

The system of differential equations was solved numerically using the scipy odeint library (python 3.8) with initial conditions specified in **Table 2**.

**Table 2**. Values and descriptions of initial conditions that were used to solve the system of differential equations.

| Variable | Value | Description |
|---|---|---|
| N(t=0) | 1.53e10 CFU/mL | Maximal average growth capacity estimated from optical density of *B. fragilis* cultures – CPS 'phase-locked' mutants and phage resistant-isolates – in the absence of phage |
| $B_{Sus}$(t=0) | 8.75e8 CFU/mL,<br>8.75e8 CFU/mL,<br>8.75e8 CFU/mL,<br>8.02e8 CFU/mL,<br>7.29e9 CFU/mL,<br>2.92e8 CFU/mL | Initial concentration of phage-sensitive cells that represent, respectively, 0.6 (for $B_{Fp}$(t=0) 0.005, 0.01, and 0.015), 0.55, 0.50, and 0.20 fractions of all cells at time point 0. Estimated based on the initial optical density of the experimental *B. fragilis* culture |
| $B_{Fp}$(t=0) | 7.29e6 CFU/mL,<br>1.46e7 CFU/mL,<br>2.19e7 CFU/mL,<br>7.29e7 CFU/mL,<br>1.46e8 CFU/mL,<br>5.83e8 CFU/mL | Tested initial concentrations of phage 'super-resistant' cells which represent, respectively, 0.005, 0.01, 0.015, 0.05, 0.1, and 0.4 fractions of all cells at time point 0 |
| P(t=0) | 87500 PFU/mL | Initial concentration of the Bf12P1 phage. Concentration is known because a phage aliquot with a known titer was used in the experiment |

Modeling results are shown in **Supplementary Figures 23E-F**. The model captures the main growth stages of the *wt B. fragilis* culture in the presence of Bf12P1, such as initial growth of phage-sensitive cells (up to 4 h) followed by culture collapse (at 4 h) caused by dramatic lysis, and a slow re-growth of phage-resistant cells after 10-12 h (**Supplementary Figure 23E**). Varying initial concentrations of 'super-resistant' cells, we showed that those cells likely comprise a fraction between 0.5% and 1.5% of the initial *wt* population (**Supplementary Figure 23F**). Modelling of increased fractions of 'super-resistant' cells showed earlier outgrowth of a culture and a different overall shape of a growth curve (for 40% fraction) (**Supplementary Figure 23E**).

It should be noted that the model has several limitations. First, parameters were fitted based on optical density data, which is not a perfect proxy for the concentration of viable cells. Second, upon lysis, cell debris creates residual optical density which cannot be described by the model, causing deviations from the experimental data (see the region between 5 and 16 h in the plots). Third, the model does not account for phage absorption on dead and 'super-resistant' cells which might cause a reduction in the effective phage concentration. Fourth, the model does not account for possible switching between phage-sensitive and 'super-resistant' categories during the culture growth caused by ongoing phase variation or epigenetics. Particularly, we speculate that the seemingly reduced growth rates of ‘super-resistant’ cells at late stages of growth (after 20 h), which is not currently shown by the model, may be caused by switching to phage-sensitive cells.

### Supplementary Figures

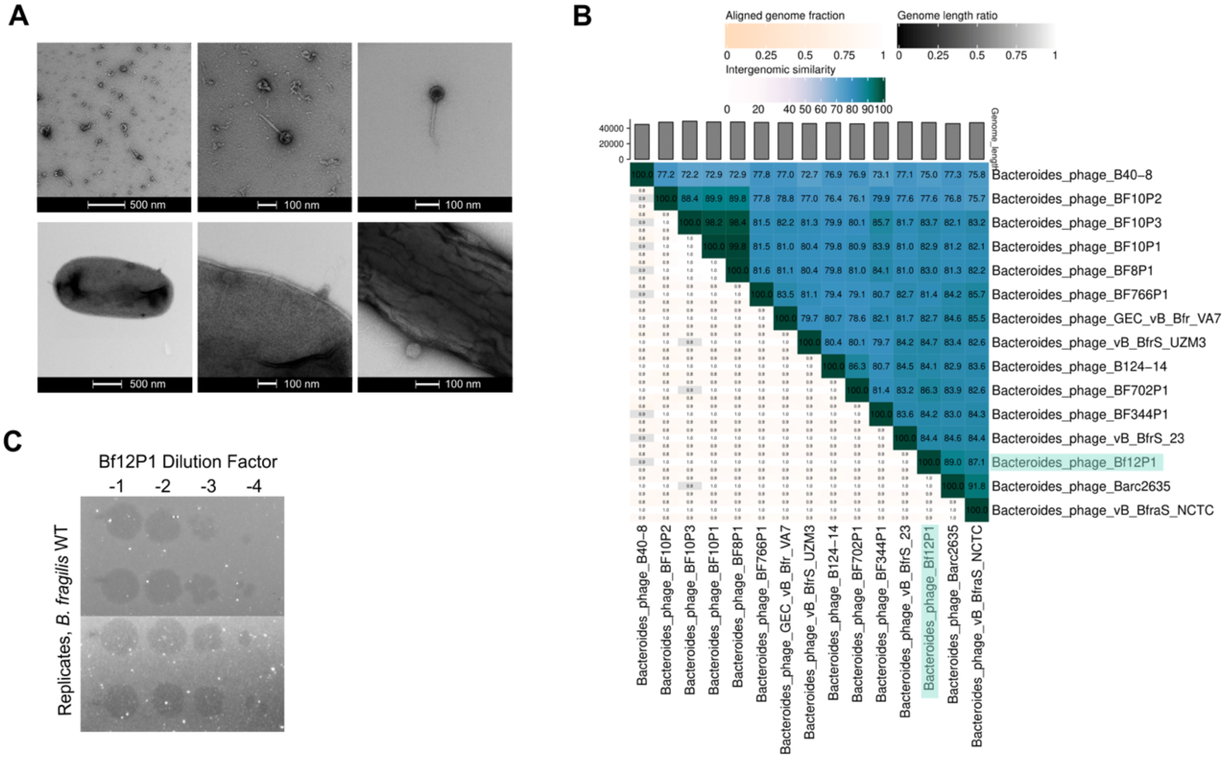


**Supplementary Figure 1**

**Characterization of a novel lytic bacteriophage Bf12P1. (A)** Transmission electron microscopy images of Bf12P1 phage alone and with *B. fragilis*. **(B)** Comparison of Bf12P1genome to those of closely related viruses indicating intergenomic similarity, performed by VIRIDIC. The most closely related genome to Bf12P2 is Bacteroides phage Barc2635 with 89.0% similarity. **(C)** Visible lysis of *B. fragilis* on plates with Bf12P1, in different concentrations as indicated by dilution factors.

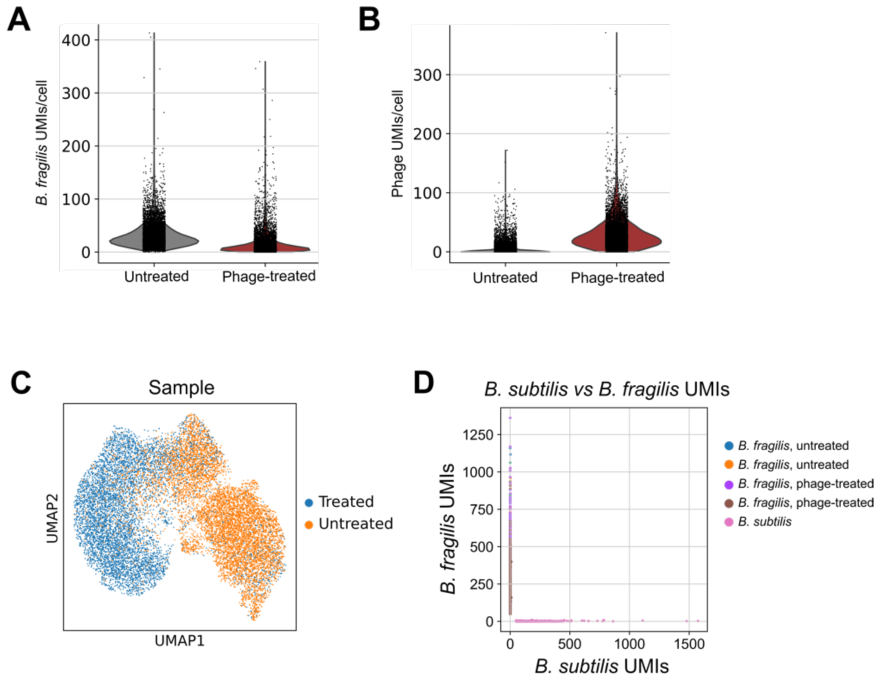


**Supplementary Figure 2**

**M30 experiment performed prior to F3 yielded slightly lower mRNA capture but similar overall findings to F3. (A)** *B. fragilis*, and **(B)** Bf12P1 filtered mRNA UMIs/cell for untreated and phage-treated samples from M30 experiment. **(C)** UMAP embedding of both treated and untreated samples from M30, using both *B. fragilis* and Bf12P1 genes, colored by sample. **(D)** Alignment of M30 cells to concatenated *B. fragilis* and *B. subtilis* genomes demonstrates single-cell resolution of the experiment.

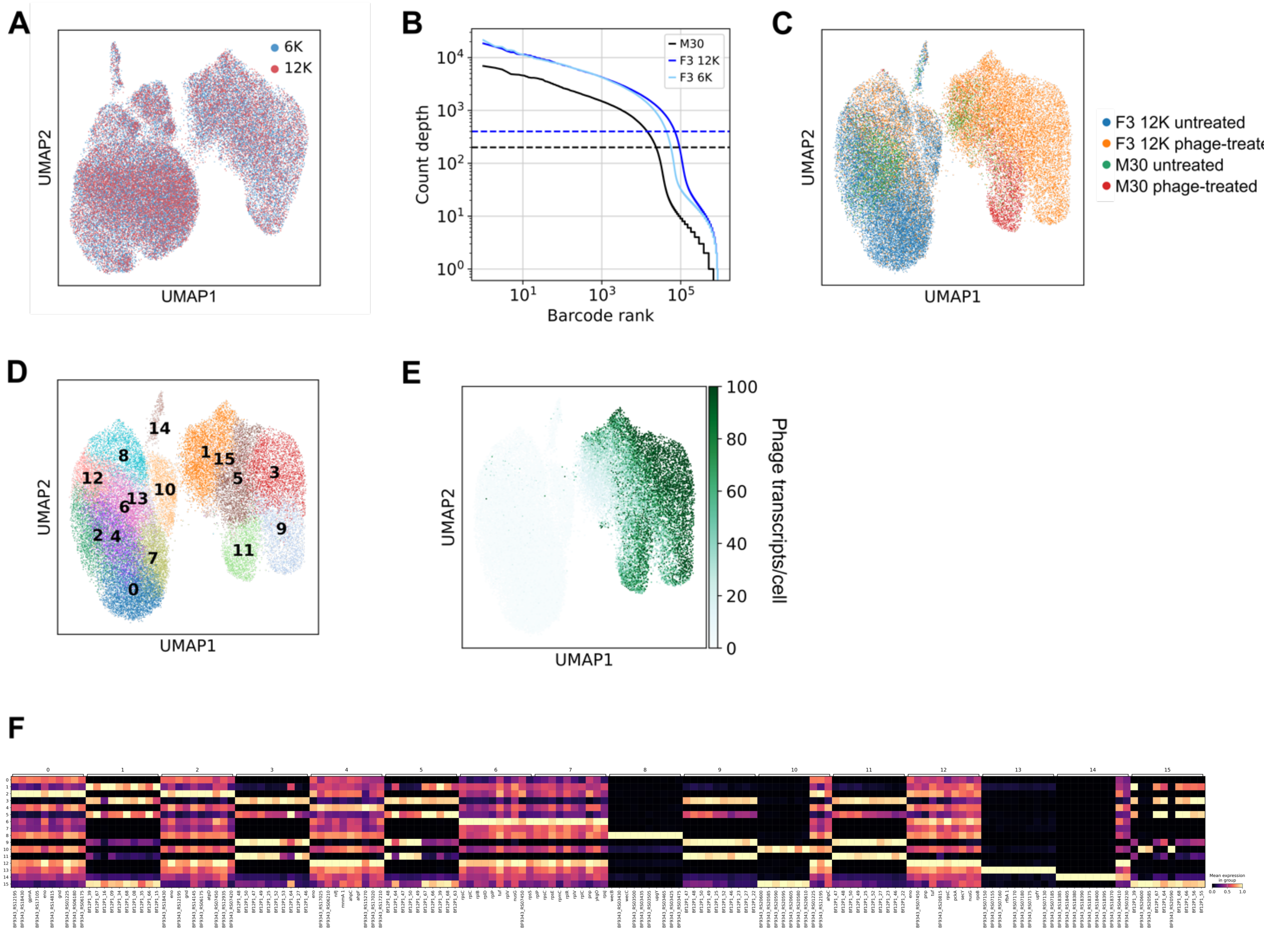


**Supplementary Figure 3**

**Comparison of data from M30 and two replicate libraries of F3 experiments. (A)** UMAP embedding of all samples, using both *B. fragilis* and Bf12P1 genes, colored by technical replicate libraries from the F3 experiment (labeled 6K and 12K). **(B)** Comparison of initial UMI/cell filtering thresholds for M30 experiment and both libraries from the F3 experiment, showing that F3 experiment yielded higher resolution data. **(C-E)** UMAPs of combined phage-treated and untreated samples for F3 12K library and M30, colored by either sample from each experiment (C), clusters by Louvain clustering algorithm (D), or abundance of phage transcripts per cell (E). **(F)** Top 10 expressed genes corresponding to each Louvain cluster from panel D.

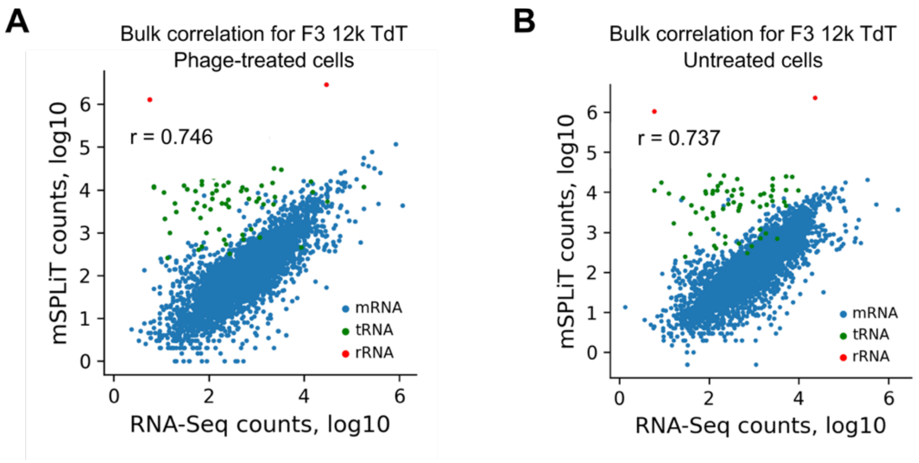


**Supplementary Figure 4**

**Correlation of expression between summed microSPLiT data and bulk RNA sequencing.** Correlation (Pearson) of expression between summed expression data from (A) phage-treated cells, and (B) untreated cells in the F3 12K library compared to bulk RNA sequencing of phage-treated sample. Genes with zero expression levels were excluded, while tRNA/rRNA were included.

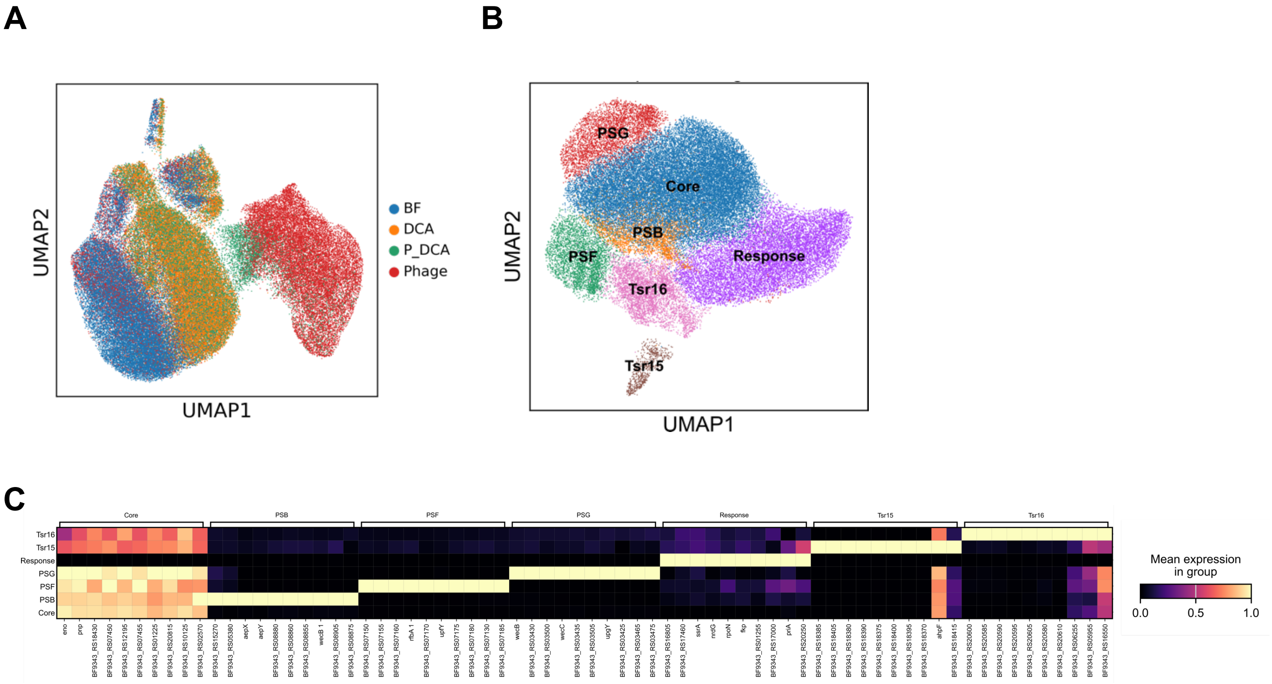


**Supplementary Figure 5**

**Clusters identified in microSPLiT dataset. (A)** UMAP of all samples with *B. fragilis* and phage genes, colored by sample name: untreated (BF), phage-treated (Phage), DCA-treated (DCA), and both phage and DCA treatments (P_DCA). **(B)** UMAP of the untreated and phage-treated samples, using only *B. fragilis* genes from the no-DCA dataset, colored by labeled Louvain clusters and **(C)** corresponding top 10 genes for each cluster.

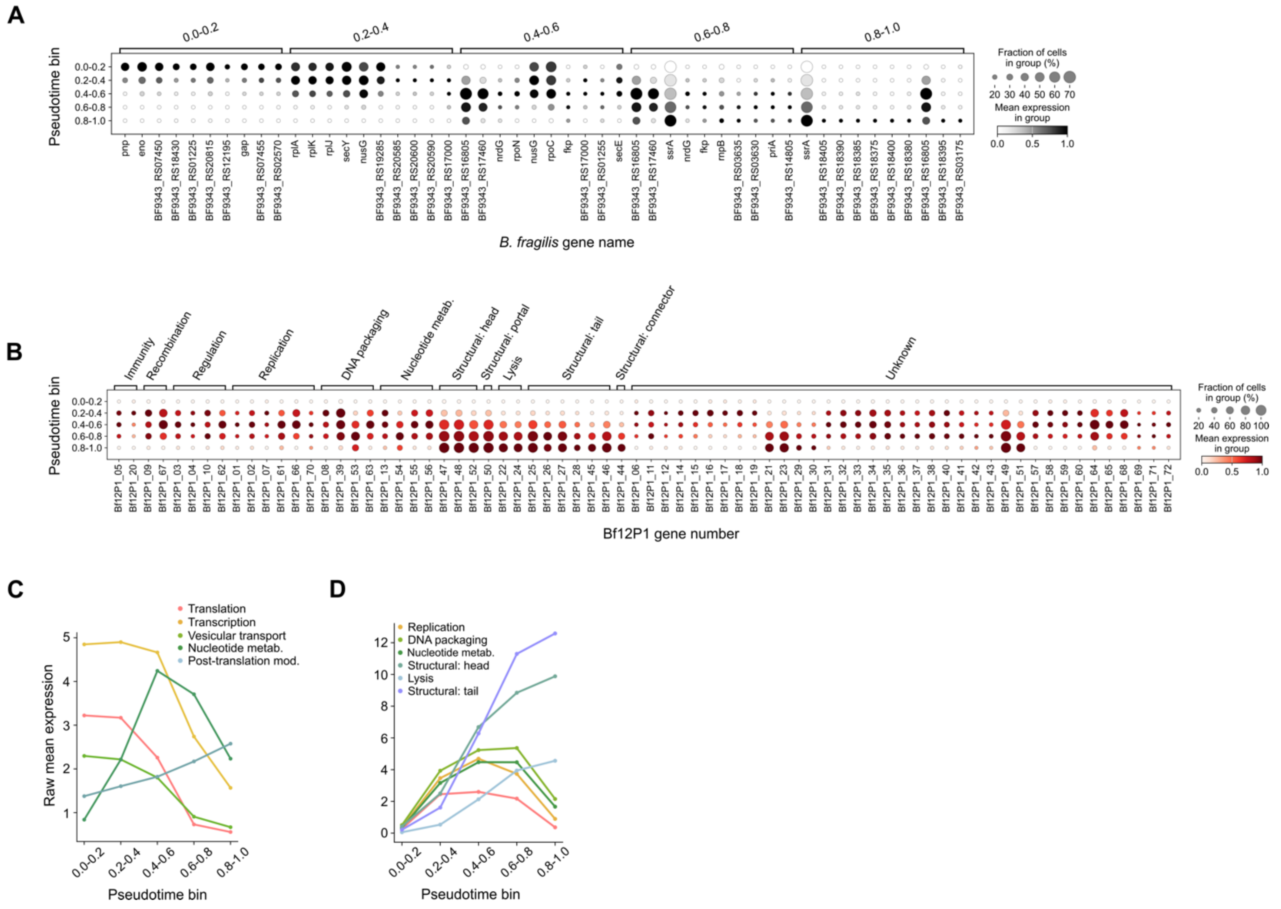


**Supplementary Figure 6**

**Patterns of both *B. fragilis* and phage gene expression along pseudotime. (A)** Top 10 *B. fragilis* genes expressed by cells categorized into each pseudotime bin, colored by the normalized expression, and used for subsequent functional category analysis. **(B)** All Bf12P1 genes, colored by the normalized expression in each pseudotime bin and sorted into functional categories, including genes with unknown functions. **(C-D)** Relative abundance of **(C)** selected *B. fragilis* functional categories and **(D)** selected Bf12P1 functional categories over pseudotime bins.

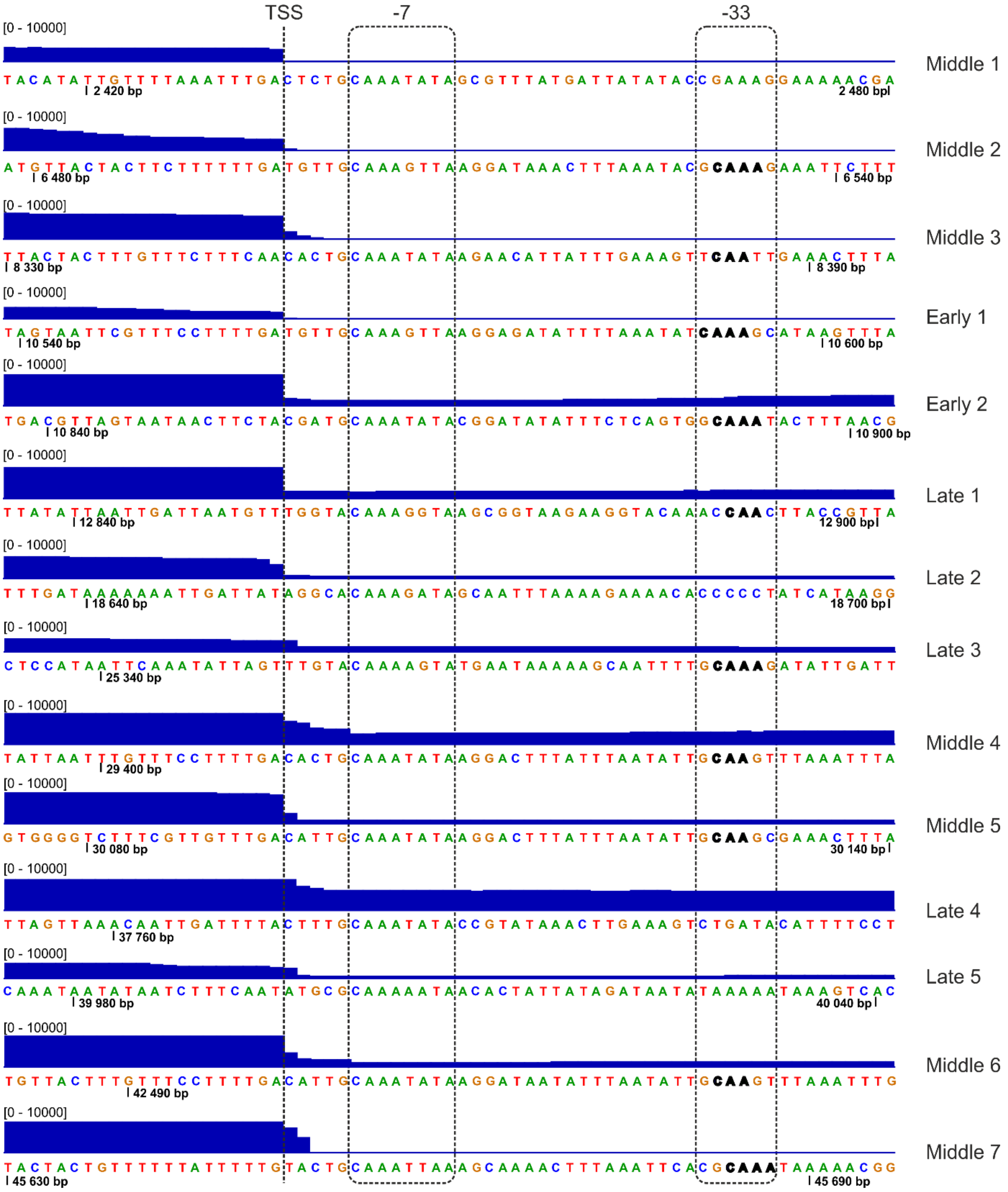


**Supplementary Figure 7**

**Transcription start sites (TSS) and promoters identified in the Bf12P1 genome.** MicroSPLiT read density (blue track) and sequence are shown for each putative TSS. Positions of TSS, -7 and -33 elements of putative promoters are shown with dashed lines and rectangles.

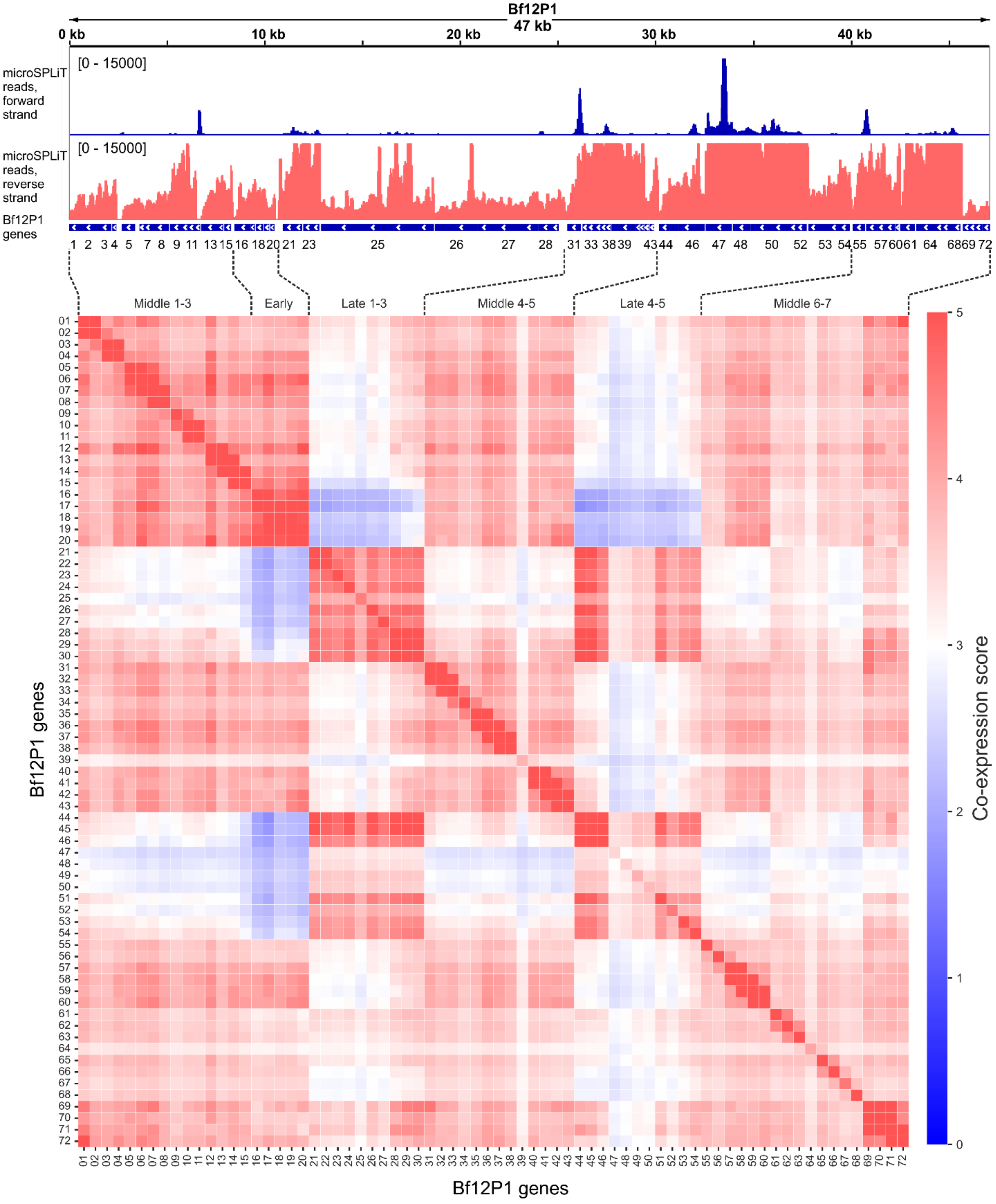


**Supplementary Figure 8**

**Co-expression of Bf12P1 genes.** Density of microSPLiT reads for the Bf12P1 genome is shown on top for forward (top track, blue) and reverse (a track below, red) strands. Coverage depth limits are indicated in brackets. The heatmap below represents co-expression scores calculated between

Bf12P1 genes. Gene clusters identified based on the following: a) reads density, b) clusters of co-expressed genes, c) median pseudotime values (**Figure 2K**), are labeled between genomic tracks and the heatmap.

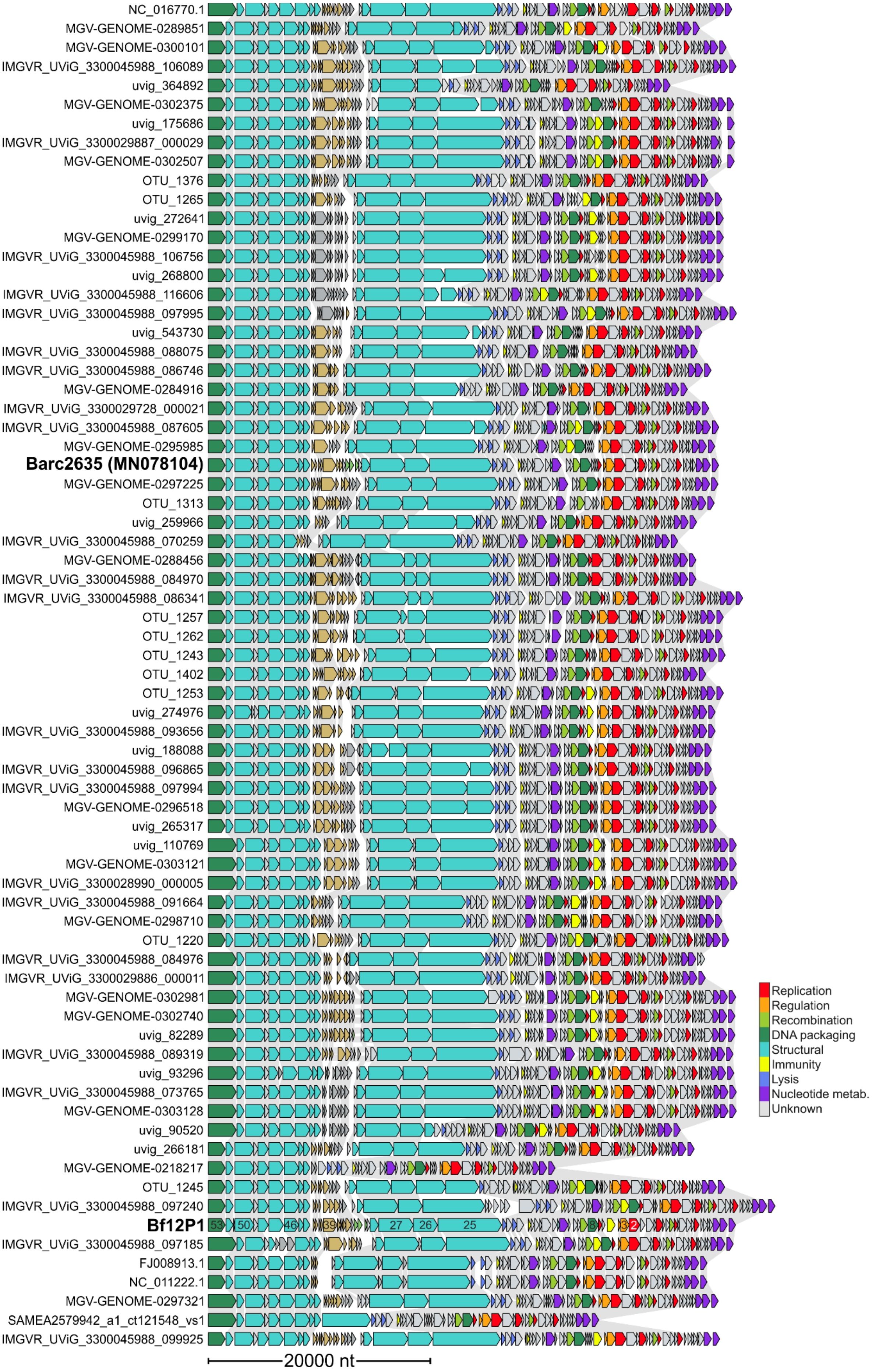
NC_016770.1
MGV-GENOME-0289851
MGV-GENOME-0300101
IMGVR_UViG_3300045988_106089
uvig_364892
MGV-GENOME-0302375
uvig_175686
IMGVR_UViG_3300029887_000029
MGV-GENOME-0302507
OTU_1376
OTU_1265
uvig_272641
MGV-GENOME-0299170
IMGVR_UViG_3300045988_106756
uvig_268800
IMGVR_UViG_3300045988_116606
IMGVR_UViG_3300045988_097995
uvig_543730
IMGVR_UViG_3300045988_088075
IMGVR_UViG_3300045988_086746
MGV-GENOME-0284916
IMGVR_UViG_3300029728_000021
IMGVR_UViG_3300045988_087605
MGV-GENOME-0295985
Barc2635 (MN078104)
MGV-GENOME-0297225
OTU_1313
uvig_259966
IMGVR_UViG_3300045988_070259
MGV-GENOME-0288456
IMGVR_UViG_3300045988_084970
IMGVR_UViG_3300045988_086341
OTU_1257
OTU_1262
OTU_1243
OTU_1402
OTU_1253
uvig_274976
IMGVR_UViG_3300045988_093656
uvig_188088
IMGVR_UViG_3300045988_096865
IMGVR_UViG_3300045988_097994
MGV-GENOME-0296518
uvig_265317
uvig_110769
MGV-GENOME-0303121
IMGVR_UViG_3300028990_000005
IMGVR_UViG_3300045988_091664
MGV-GENOME-0298710
OTU_1220
IMGVR_UViG_3300045988_084976
IMGVR_UViG_3300029886_000011
MGV-GENOME-0302981
MGV-GENOME-0302740
uvig_82289
IMGVR_UViG_3300045988_089319
uvig_93296
IMGVR_UViG_3300045988_073765
MGV-GENOME-0303128
uvig_90520
uvig_266181
MGV-GENOME-0218217
OTU_1245
IMGVR_UViG_3300045988_097240
Bf12P1
IMGVR_UViG_3300045988_097185
FJ008913.1
NC_011222.1
MGV-GENOME-0297321
SAMEA2579942_a1_ct121548_vs1
IMGVR_UViG_3300045988_099925
20000 nt
Replication
Regulation
Recombination
DNA packaging
Structural
Immunity
Lysis
Nucleotide metab.
Unknown

**Supplementary Figure 9**

**Genomic comparison of Bf12P1, Barc2635, and related phages from the "proteome community" 506** (*2*)**.** Genes are color-coded according to their predicted functions. Genes of a variability hotspot corresponding to a cluster of Bf12P1 middle genes (genes 29-43) are colored in light brown.

A

| Gene (old ID) | Gene (new ID) | Predicted function | Signal peptide |
|---|---|---|---|
| 4127 | RS20620 | Tyrosine site-specific recombinase (Tsr16) | - |
| 4126 | RS20615 | AraC-family transcription factor | - |
| 4124 | RS20610 | Porin-like protein | + (Sec/SPI) |
| 4123 | RS20605 | Contains tetratricopeptide repeat | + (Sec/SPII) |
| 4122 | RS20600 | Structural similarity to major **fimbrial** subunit protein | + (non-canonical) |
| 4121 | RS20595 | Putative FimB/Mfa2 family **fimbrial** subunit | + (non-canonical) |
| 4120 | RS20590 | Structural similarity to minor **fimbrium** tip subunit protein mfa3 | + (Sec/SPII) |
| 4119 | RS20585 | Contains BACON domain | + (Sec/SPII) |
| hup4 | RS20580 | HU-like protein | - |

B

| # of cells | Tsr16 gene cluster not expressed | Tsr16 gene cluster expressed |
|---|---|---|
| 4125-26 region ref. orientation | 76 | 12 |
| 4125-26 region inverted | 120 | 284 |

Fisher's exact test
p-value < 0.0001

**Supplementary Figure 10**

**The structure and transcription regulation of the Tsr16-associated gene cluster. (A)** Functional gene of microSPLiT reads for the Tsr16-associated gene cluster. **(B)** A contingency table showing association between orientations of the regulatory region (derived from read alignment) and transcription of the Tsr16-associated gene cluster (derived from microSPLiT data).

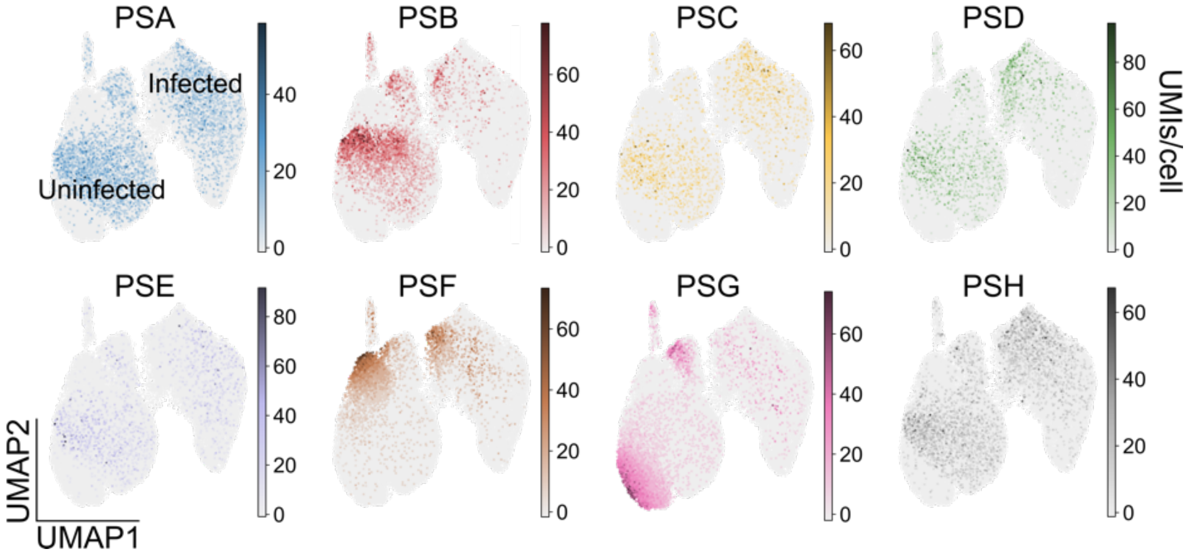


**Supplementary Figure 11**
**Distributions of cells expressing each CPS type.** Summed expression values for each of the CPS operons PSA through PSH shown on a UMAP embedding of all samples with both *B. fragilis* and phage transcripts.

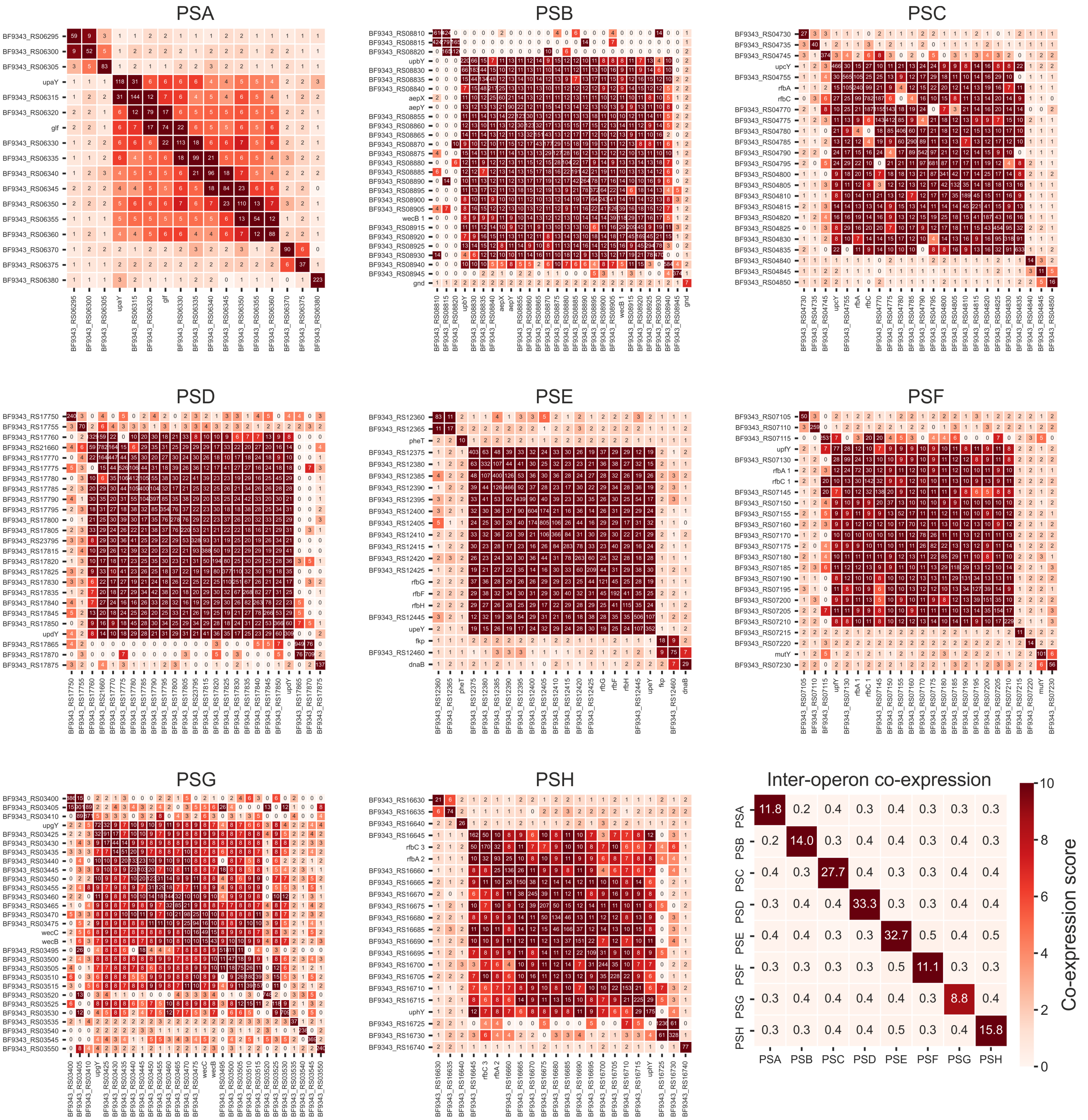


**Supplementary Figure 12**

**CPS operon and gene co-expression.** Co-expression score matrices of all CPS operons with one another (inter-operon co-expression), as well as genes within each CPS operon and 2-3 adjacent genes together (individual CPS operons).

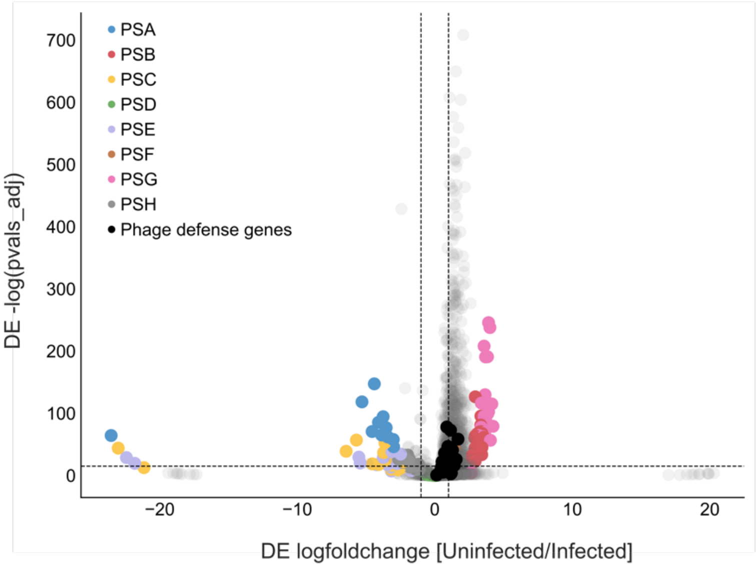


**Supplementary Figure 13**

**Differentially expressed genes between the phage-treated uninfected and infected cells.** Defense genes are genes involved in anti-bacteriophage defense systems that were annotated in the data using DefenseFinder (Supplementary Table 7). Each gene in a CPS operon is colored according to the CPS type (Supplementary Table 2).

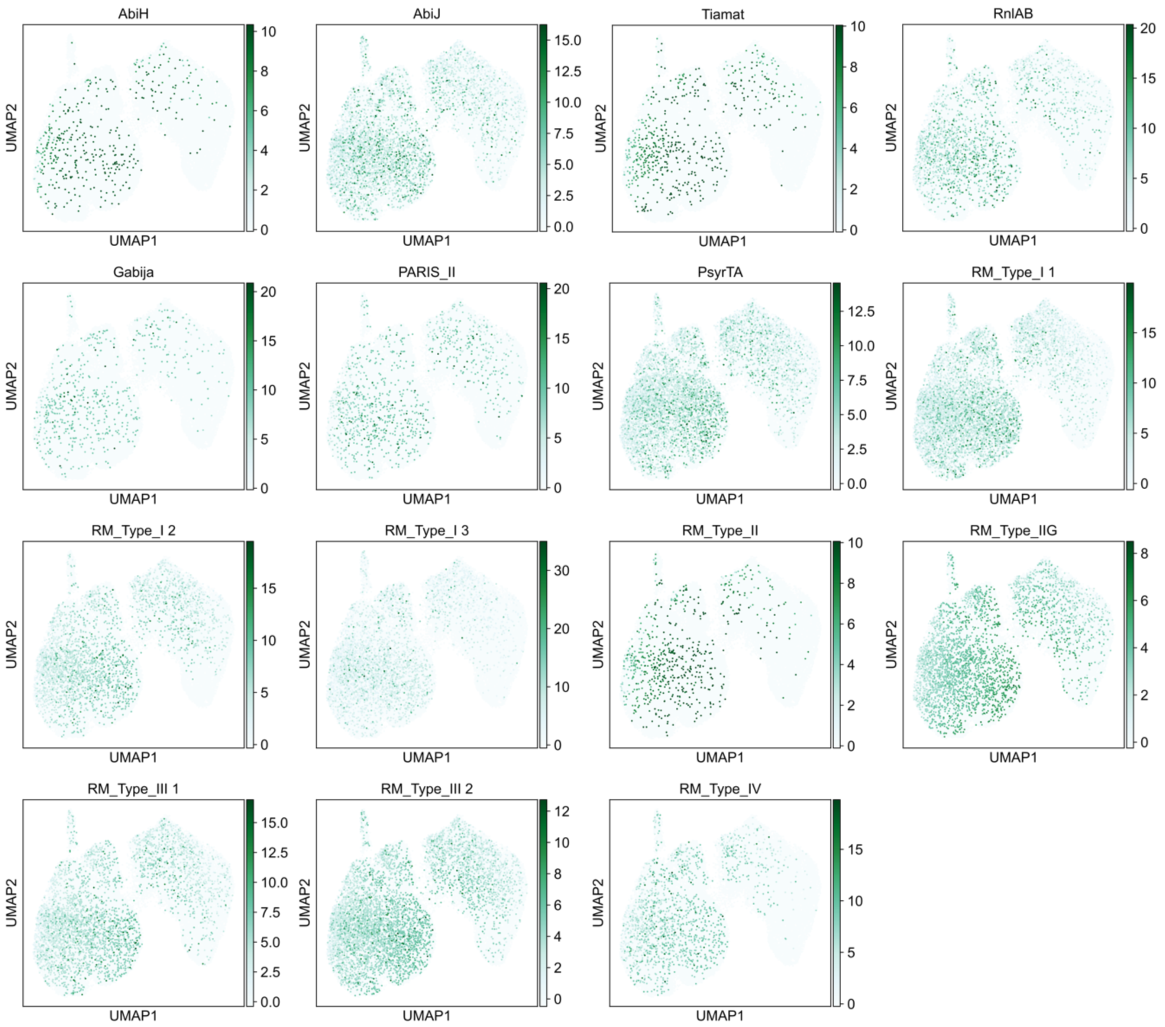


**Supplementary Figure 14**
**Putative defense systems expressed in the microSPLiT dataset.** Summed expression values for each of the defense systems, as annotated by DefenseFinder, shown on a UMAP embedding of all samples with both *B. fragilis* and phage transcripts.

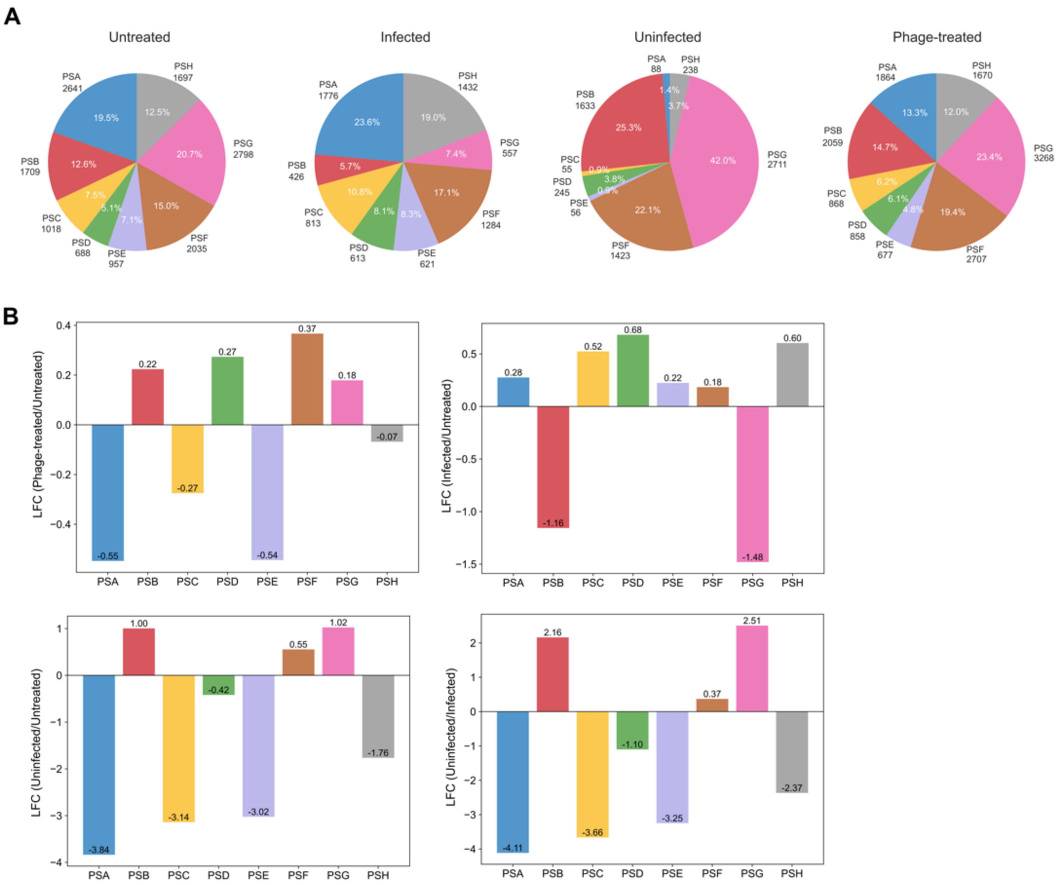


**Supplementary Figure 15**
**Distributions of cells expressing each CPS type between sample subsets. (A)** Proportions of CPS types for each sample, including untreated, phage-treated, uninfected, and infected cells. **(B)** Relative changes in fractions of each CPS type between marked subsets of cells.

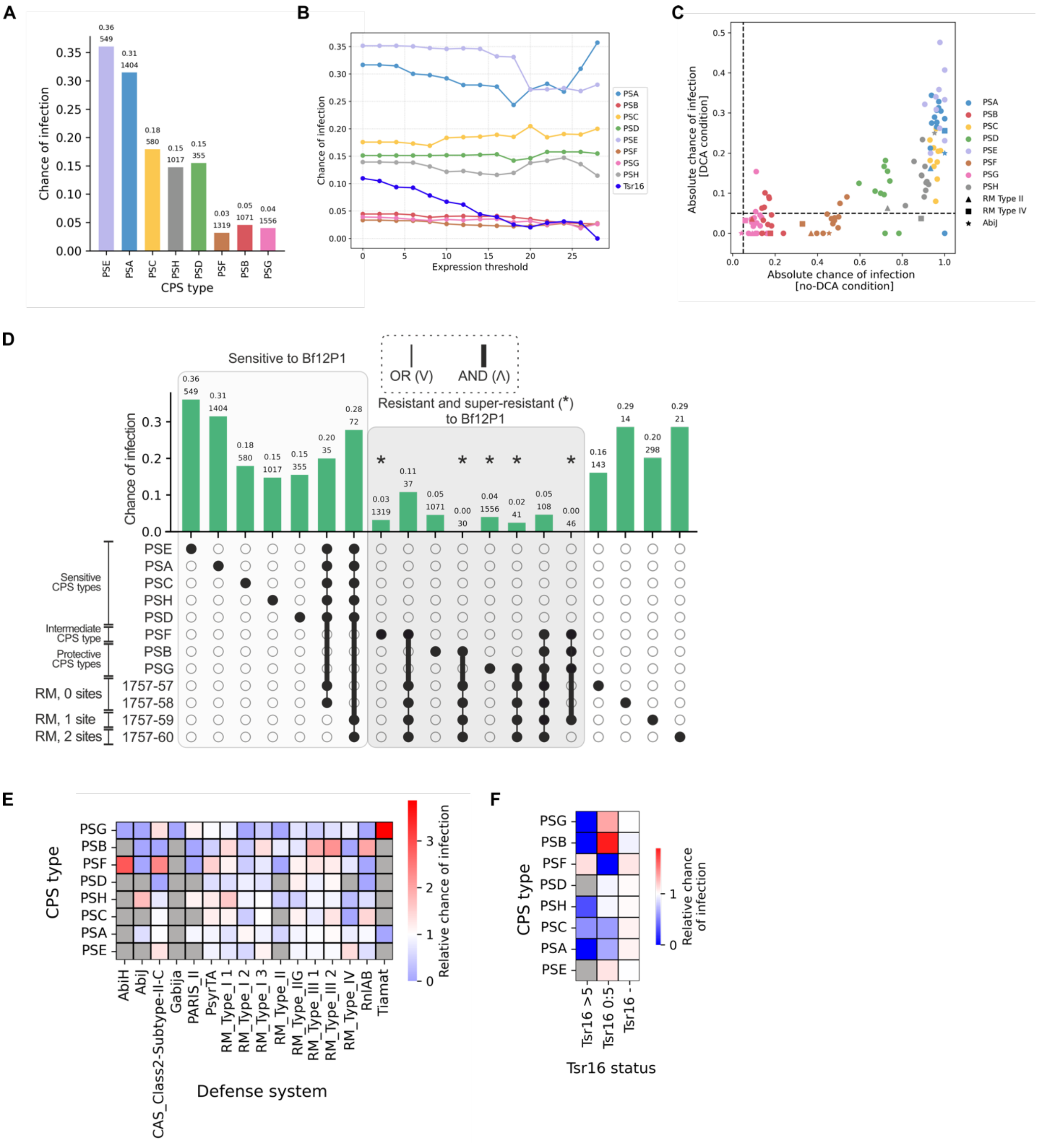


**Supplementary Figure 16**

**Comparison of infection chances for distinct phenotypes in the DCA dataset to the cells in no-DCA dataset. (A)** Infection chance for cells expressing each CPS type in DCA conditions. **(B)** Chance of infection for cells expressing each CPS type or Tsr16-associated gene cluster above multiple indicated thresholds of expression. **(C)** Correlation between absolute chance of infection values for cells expressing different CPS and defense system combinations, for the DCA and no-DCA conditions. Data is shown for combinations detected in both conditions. **(D)** Phage infection chances estimated for indicated cell subpopulations based on microSPLiT data for samples grown with addition of DCA. Infection chance and number of cells in a subpopulation are shown above each bar. Subpopulations are grouped according to the level of protection against the phage. **(E)** Relative chance of infection for DCA-treated cells co-expressing each CPS type and the indicated anti-phage defense mechanisms simultaneously. Subpopulations with less than 10 cells were excluded from the analysis (labeled in gray). **(F)** Relative chance of infection for DCA-treated cells co-expressing each CPS type and Tsr16-adjacent region above indicated expression thresholds.

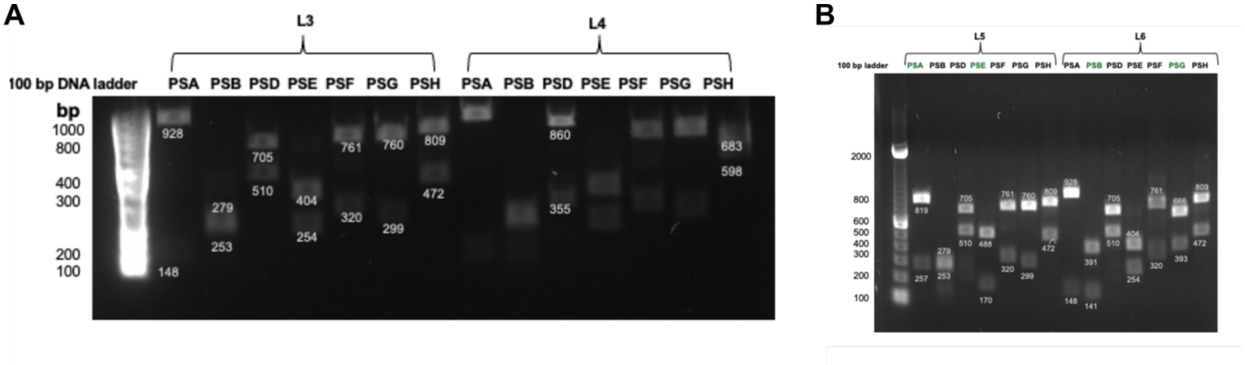


**Supplementary Figure 17**

**Verification of promoter orientations for CPS phase-locked *B. fragilis* strains. (A)** Agarose gel electrophoresis of digested amplicons from 'PSC' (L3) and "PSD/PSH" (L4) 'phase-locked' strains after PCR amplification using CPS-specific primers(*3*), confirming all expected promoter orientations. **(B)** Same as (A) for **"**PSA/PSE" (L5) and "PSB/PSG" (L6) 'phase-locked' strains.

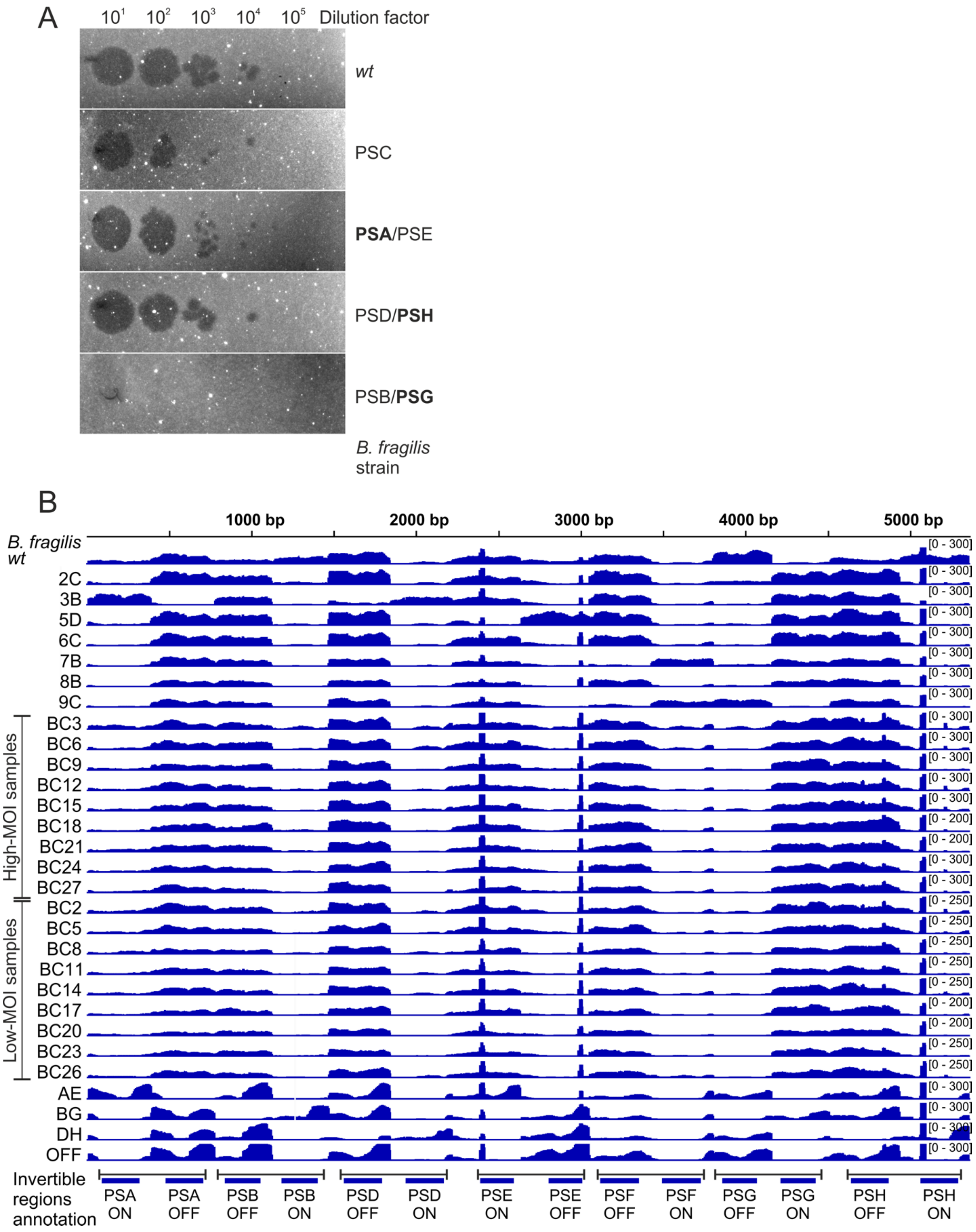


**Supplementary Figure 18**

**Phage-susceptibility of CPS 'phase-locked' strains and CPS promoters' orientation in Bf12P1-resistant isolates, bulk culture samples, and CPS-locked mutants. (A)** Representative images of phage plaque assay performed with CPS 'phase-locked' *B. fragilis* strains and the wild-type (*wt) B. fragilis* strain. Phage dilutions are indicated above the images. **(B)** IGV snapshots of read coverage depth for *wt B. fragilis*, Bf12P1-resistant isolates, and bulk culture aliquots obtained by alignment of WGS reads to the synthetic reference sequence prepared by concatenation of CPS operons promoter regions in 'OFF' and 'ON' states. The coverage depth was used for quantification of CPS operons' promoter states shown in **Figure 4D**.

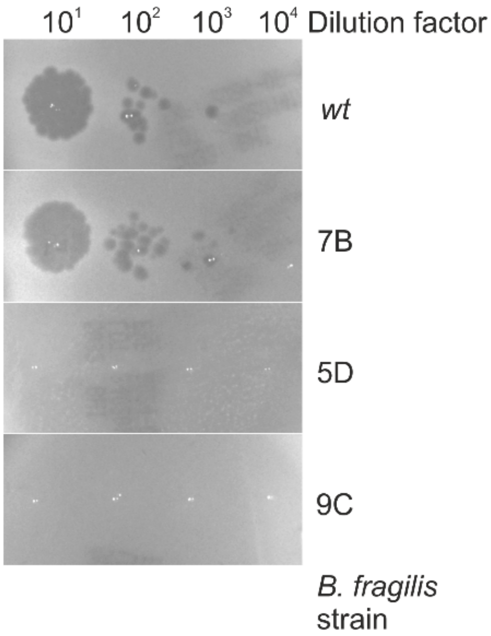


**Supplementary Figure 19**
***B. fragilis* isolates phage susceptibility.** Representative images of phage plaque assay performed with Bf12P1-resistant *B. fragilis* isolates and the *wt B. fragilis* strain. Phage dilutions are indicated above the images.

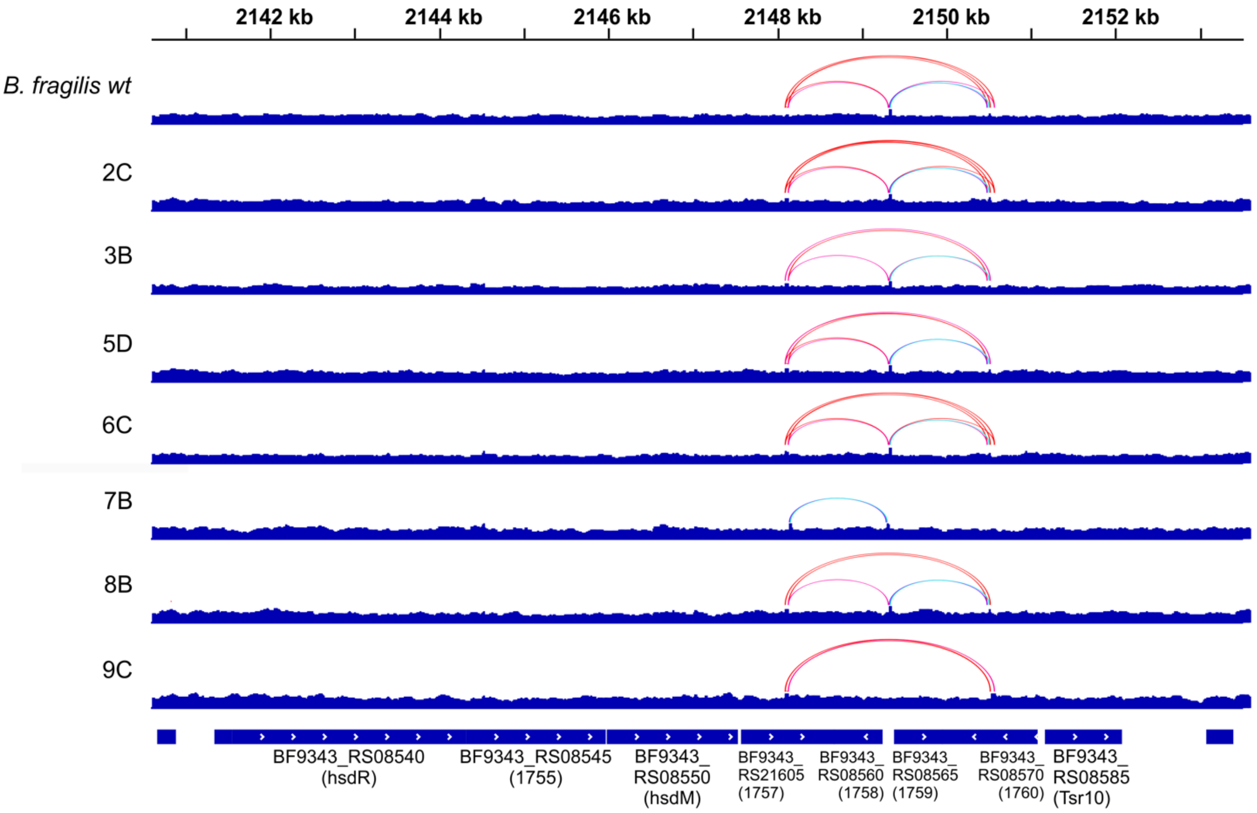


**Supplementary Figure 20**

**Detection of recombination within the phase-variable RM type-I gene cluster in the *B. fragilis* genome.** IGV snapshots of read coverage depth in the phase-variable RM type-I gene cluster for *wt B. fragilis* and Bf12P1-resistant isolates. Coverage depth was obtained by alignment of WGS reads to the NCTC9343 reference genome. Recombination events were detected by analysis of read alignments. Pairs of joint recombination sites are shown as arches.

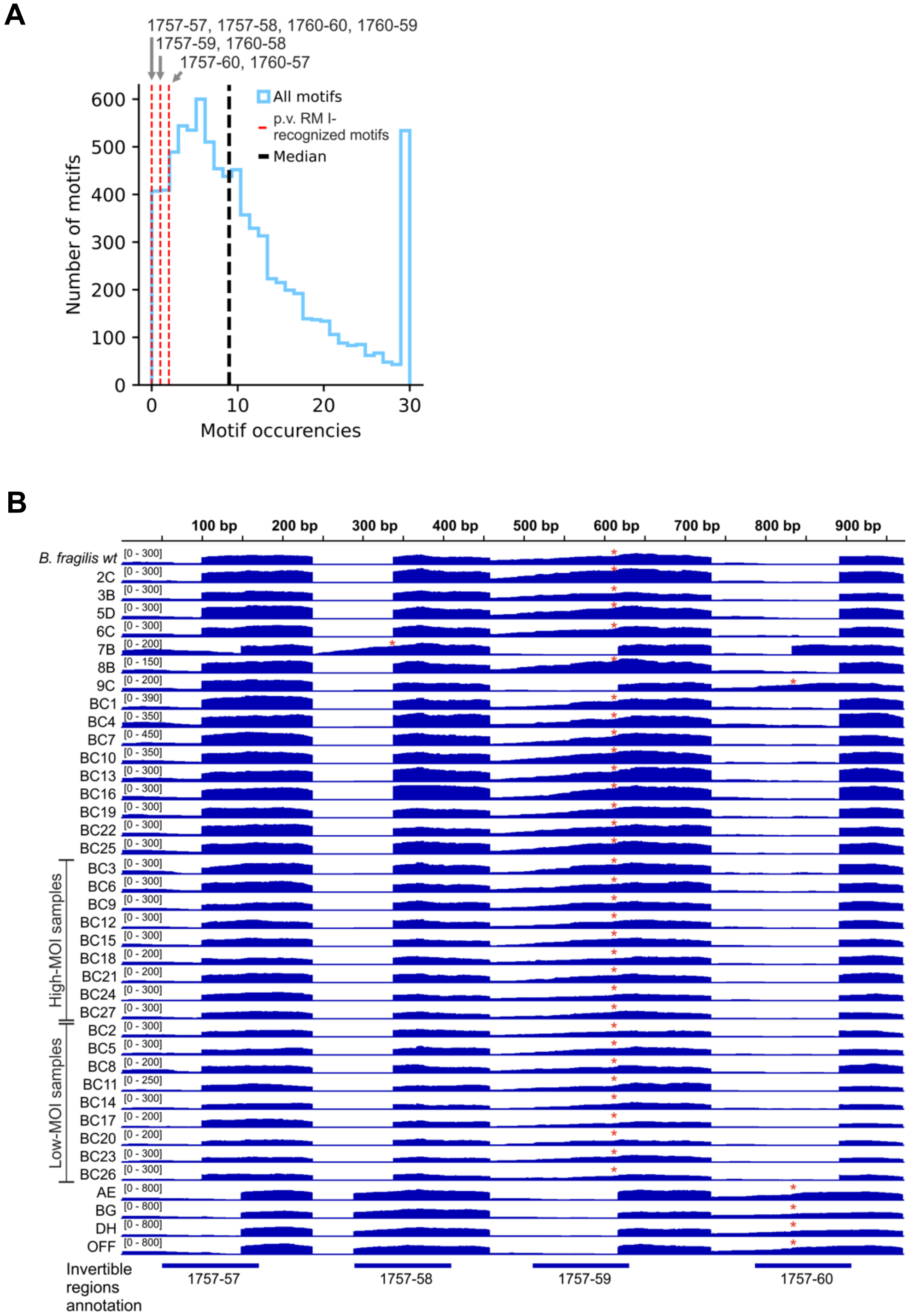


**Supplementary Figure 21**

**Phase-variable RM type-I of *B. fragilis*: numbers of sites, composition of specificity subunits, and phage susceptibility of RM type-I 'phase-locked' strains. (A)** Occurrence frequencies of 6-nucleotide motifs with similar structure and complexity to the motifs recognized by phase-variable RM type-I in the Bf12P1 genome. Actual occurrence frequencies for motifs recognized by different variants of phase-variable RM type-I specificity subunits are shown by vertical dashed red lines. Motif sequencies recognized by the phase-variable RM type-I were taken from (*4*) and (*5*). **(B)** IGV snapshots of read coverage depth for *wt B. fragilis*, Bf12P1-resistant isolates, and bulk culture aliquots. Coverage depth was obtained by the alignment of WGS reads to the synthetic reference sequence prepared by concatenation of four possible variants of a specificity subunit gene region that contains recombination sites and is involved in shuffling. Variants with the 1760 gene-encoded N-terminal part are not shown. Red asterisks indicate a specificity subunit variant detected in a sample. The coverage depth was used for determining the specificity subunit type shown in **Figure 5D**.

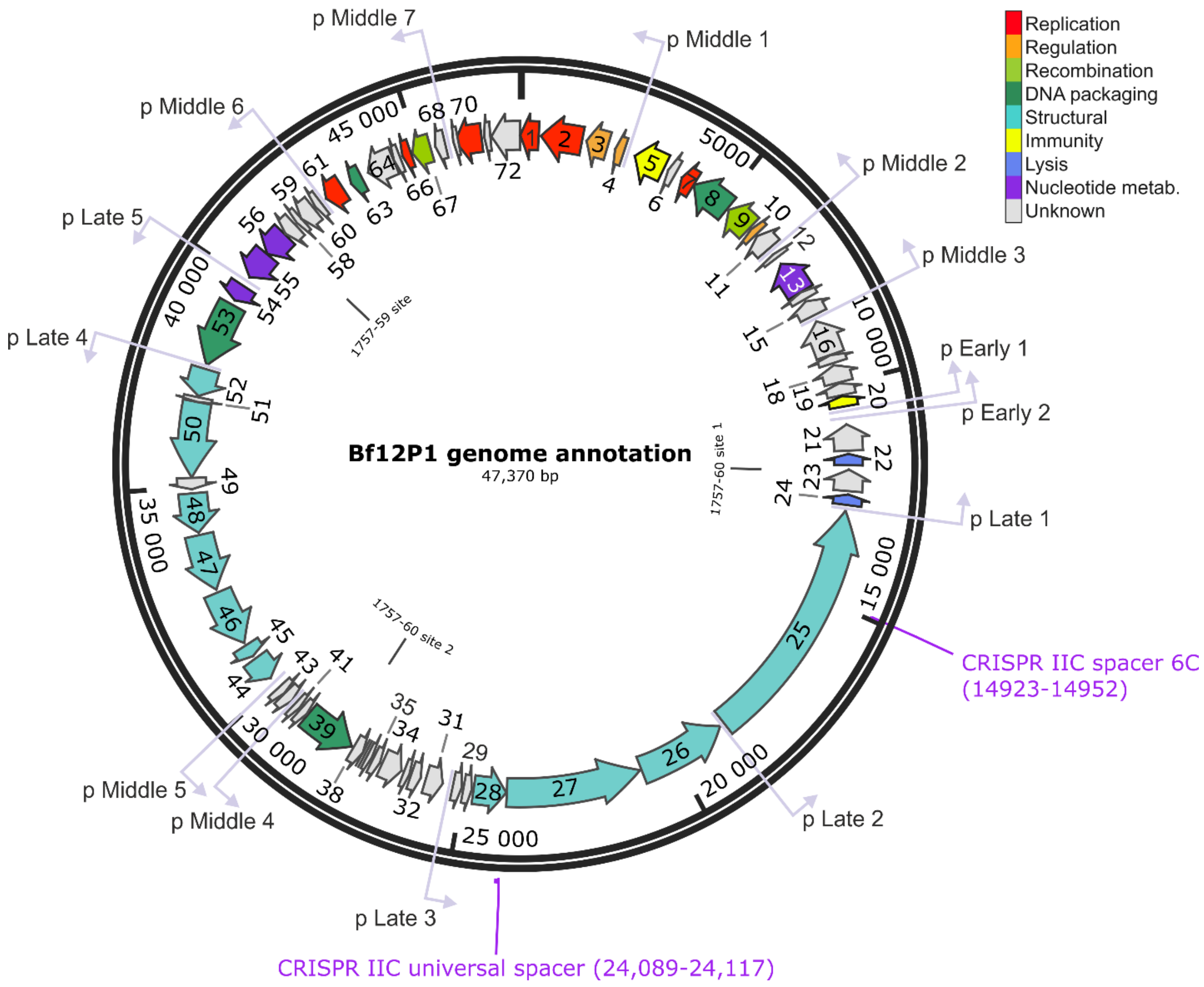


**Supplementary Figure 22**

**Location of phase-variable RM-type I cut sites and CRISPR-Cas spacers acquired by select isolates in Bf12P1 genome.** The site recognized by the 1757-59 specificity subunit is located in gene 57, while two sites recognized by the 1757-60 specificity subunit are located in genes 23 and 39. A protospacer perfectly matching the CRISPR-Cas IIC spacer acquired by the resistant isolate 6C is located in gene 25, while a degenerate protospacer with two mismatches is located in gene 28.

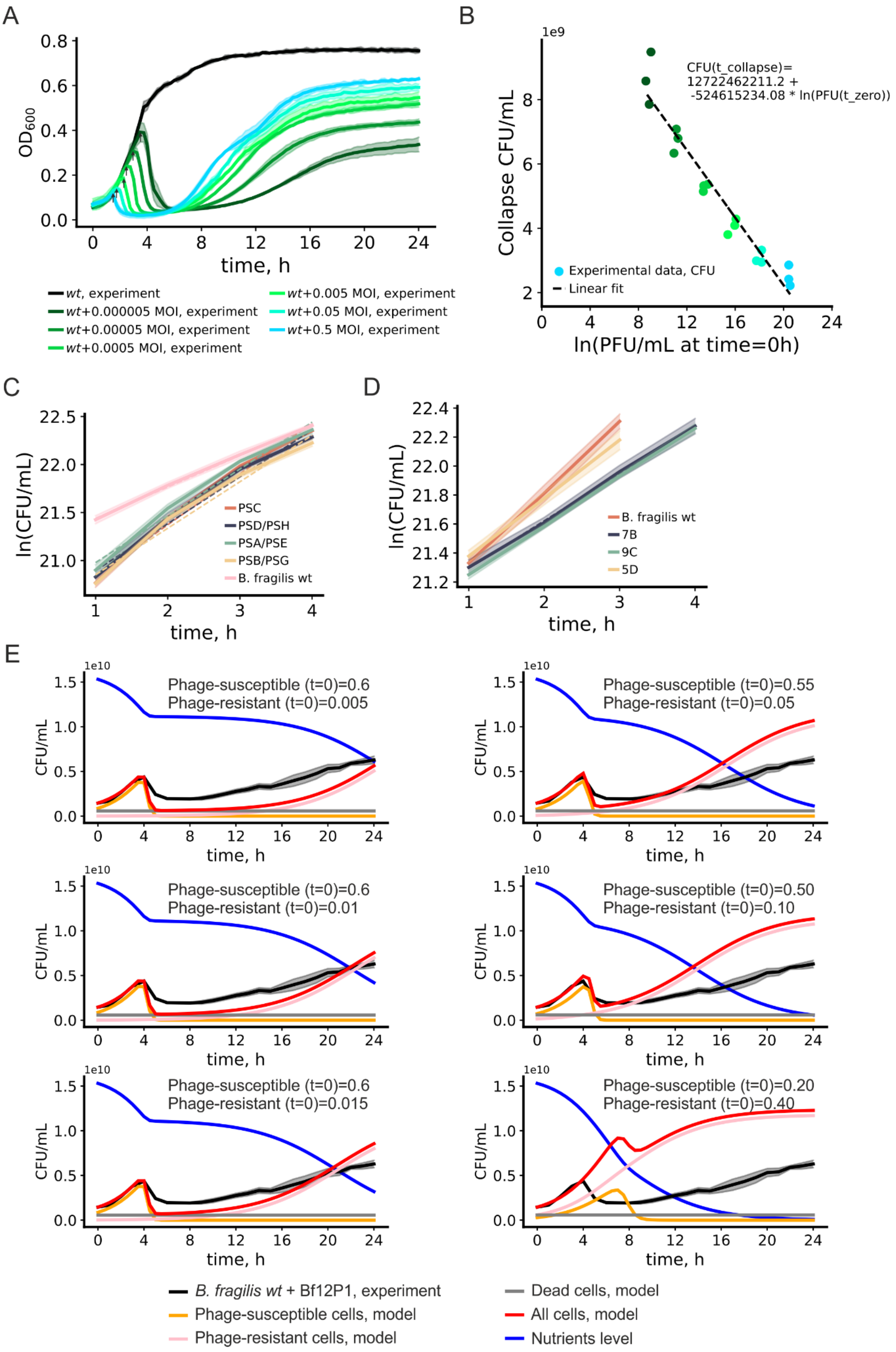
A
OD600
time, h
wt, experiment
wt+0.000005 MOI, experiment
wt+0.00005 MOI, experiment
wt+0.0005 MOI, experiment
wt+0.005 MOI, experiment
wt+0.05 MOI, experiment
wt+0.5 MOI, experiment
B
1e9
Collapse CFU/mL
CFU(t_collapse)=
12722462211.2 +
-524615234.08 * ln(PFU(t_zero))
Experimental data, CFU
Linear fit
ln(PFU/mL at time=0h)
C
ln(CFU/mL)
time, h
PSC
PSD/PSH
PSA/PSE
PSB/PSG
B. fragilis wt
D
ln(CFU/mL)
time, h
B. fragilis wt
7B
9C
5D
E
1e10
CFU/mL
time, h
Phage-susceptible (t=0)=0.6
Phage-resistant (t=0)=0.005
Phage-susceptible (t=0)=0.55
Phage-resistant (t=0)=0.05
Phage-susceptible (t=0)=0.6
Phage-resistant (t=0)=0.01
Phage-susceptible (t=0)=0.50
Phage-resistant (t=0)=0.10
Phage-susceptible (t=0)=0.6
Phage-resistant (t=0)=0.015
Phage-susceptible (t=0)=0.20
Phage-resistant (t=0)=0.40
B. fragilis wt + Bf12P1, experiment
Phage-susceptible cells, model
Phage-resistant cells, model
Dead cells, model
All cells, model
Nutrients level

F

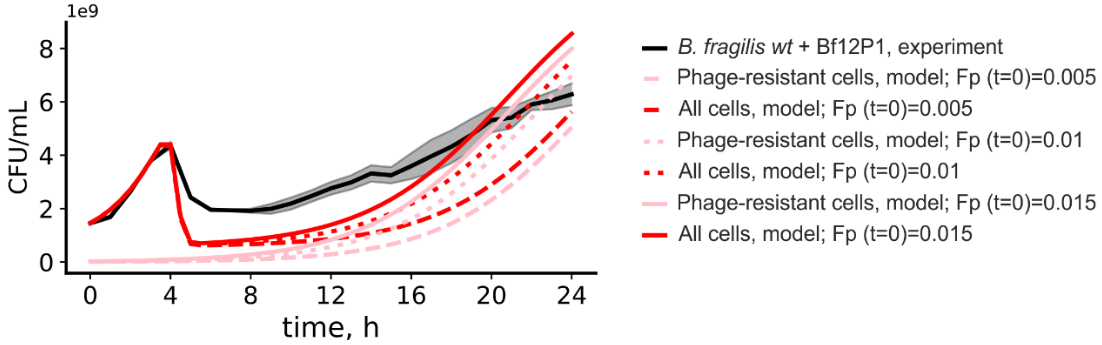


**Supplementary Figure 23**

**Modelling of *B. fragilis* culture with Bf12P1 phage. (A)** Growth curves for the liquid culture phage assay performed with *B. fragilis wt* culture. Bf12P1 was added at time point 0 at indicated multiplicities of infection (MOIs); black curve shows untreated sample. Shaded curves represent a 0.95 confidence interval of the mean of three biological replicates. Black arrows indicate growth curve points where *B. fragilis* cultures started to collapse due to lysis by Bf12P1. **(B)** *B. fragilis* culture concentration at culture collapse points (indicated in panel B) as a function of logarithm of initial Bf12P1 titer. Culture concentration in CFU/mL was assessed by optical density of the culture. Dashed line represents linear regression. **(C)** Quantification of growth rate constants for CPS 'phase-locked' *B. fragilis* strains and *wt B. fragilis*. Dashed lines represent linear regressions. Shade represents 0.95 confidence intervals of the mean of culture concentration (solid line). **(D)** Quantification of growth rate constants for phage-resistant *B. fragilis* strains and *wt B. fragilis*. Legend is the same as for panel C. **(E)** Modelling of *wt B. fragilis* culture growth in the presence of Bf12P1 phage. Black curve with shadow (the 0.95 confidence interval) shows experimental data. Orange and red curves represent modelled dynamics for, respectively, phage-sensitive and phage 'super-resistant' groups of cells. Gray curve and red curves represent non-dividing cells (at time point 0) and total concentration of cells (cells of all categories), respectively. The green curve represents concentration of Bf12P1 particles; the blue curve represents concentration of nutrients available for *B. fragilis* growth. Three plots show modelled dynamics for *B. fragilis* cultures with different initial fractions of phage 'super-resistant' cells (0.005-0.40). **(F)** Overlay of modelled dynamics for phage 'super-resistant' groups of *B. fragilis* cells and total concentrations of cells showed on panel E. Culture concentrations in CFU/mL were assessed by optical density of cultures for panels B-F.
